# Mapping the genetic architecture of human cortical expansion and its links to neuropsychiatric disorders

**DOI:** 10.64898/2026.06.22.733725

**Authors:** Matthew Rosenblatt, Marlee Vandewouw, Henry Fox-Jurkowitz, Camille M. Williams, Yuanxin Zhong, Da Zhi, Yingzhe Zhang, Margaret L. Westwater, Armin Raznahan, Michael J. Gandal, Tian Ge, Dustin Scheinost, Jordan W. Smoller, Travis T. Mallard

**Affiliations:** Center for Precision Psychiatry, Department of Psychiatry, Massachusetts General Hospital, Boston, MA, USA; Department of Psychiatry, Harvard Medical School, Massachusetts General Hospital, Boston, MA, USA; Psychiatric and Neurodevelopmental Genetics Unit, Center for Genomic Medicine, Massachusetts General Hospital, Boston, MA, USA; Stanley Center for Psychiatric Research, Broad Institute of MIT and Harvard, Boston, MA, USA; Laboratoire de Sciences Cognitives et Psycholinguistique, Département d′Études Cognitives, École Normale Supérieure, EHESS, CNRS, PSL University, Paris, France; Department of Psychology, University of Texas at Austin, Austin, TX, USA; Population Research Center, University of Texas at Austin, Austin, TX, USA; Department of Psychiatry, University of Oxford, Oxford, UK; Department of Radiology and Biomedical Imaging, Yale School of Medicine, New Haven, CT, USA; Yale Biomedical Imaging Institute, Yale School of Medicine, New Haven, CT, USA; Section on Developmental Neurogenomics, Human Genetics Branch, National Institute of Mental Health Intramural Research Program, Bethesda, MD, USA; Department of Psychiatry, Perelman School of Medicine, University of Pennsylvania, Philadelphia, PA, USA; Lifespan Brain Institute at Penn Med and the Children’s Hospital of Philadelphia, Philadelphia, PA, USA; Department of Biomedical Engineering, Yale University, New Haven, CT, USA; Interdepartmental Neuroscience Program, Yale University, New Haven, CT, USA; Child Study Center, Yale School of Medicine, New Haven, CT, USA; Department of Statistics and Data Science, Yale University, New Haven, CT, USA

## Abstract

Cortical expansion is a defining feature of human evolution and neurodevelopment, involving the tangential growth and gyrification of the cerebral cortex. Although disruptions to the expansion of the cortex are implicated in diverse brain-based disorders, the genetic architecture underlying this process remains undercharacterized. Here, we integrated GWAS data for five neuroanatomical phenotypes measured in up to 73,800 individuals to model a latent genomic factor capturing the shared genetic basis of cortical surface area, folding, curvature, gyrification, and intracranial volume. Using a multivariate framework, we mapped this pleiotropic architecture across biological scales, identifying novel effector genes, neural cell types, and developmental pathways involved in cortical expansion. Functional genomic evidence indicated that a diverse cellular ensemble contributes to the tangential growth and gyrification of the cortex during prenatal neurodevelopment, with progenitor cell lineages prominently involved. Regional analyses further revealed that genetic influences are spatially heterogeneous and organized along canonical anatomical, functional, and developmental gradients. Finally, cortical expansion exhibited substantial genetic overlap with neurodevelopmental, psychiatric, and neurological disorders, with shared genetic influences often concentrated in prefrontal cortical regions. Together, these findings map the genetic architecture of human cortical expansion and implicate it as a common etiological axis linking evolution, neurodevelopment, and health and disease.

## Introduction

The expansion of the cerebral cortex is one of the most distinctive aspects of human evolution and neurodevelopment, characterized by the tangential growth and gyrification of the cerebral mantle^1,2^. Beginning during early prenatal development and continuing through adolescence^3,4^, cortical expansion—particularly in the prefrontal cortex—supports higher-level cognition by providing the structural architecture for complex neural circuitry. Indeed, marked disruptions to this expansion occur in a range of neurogenetic disorders, in which significant cognitive difficulties, and even intellectual disability, are common^5^. Large-scale neuroimaging studies further underscore the relevance of this process, as morphological features related to expansion (e.g., surface area, gyrification, folding) have been robustly associated with cognitive abilities^6,7^ and various forms of psychopathology^8–10^. Although these findings suggest that delayed or atypical patterns of cortical expansion may represent a transdiagnostic liability, our understanding of the specific molecular pathways involved remains underdeveloped.

With the advent of large-scale imaging genomics resources, it has become clear that individual differences in morphometric indices of cortical expansion are heritable, polygenic, and regionally specific^11–13^. New genome-wide association study (GWAS) data offer a promising avenue to begin mapping the genetic architecture of this core neurodevelopmental process that affects both cognition and psychopathology. To date, most imaging GWAS work on cortical expansion has focused on surface area, a reliable index of tangential growth derived from magnetic resonance imaging (MRI), identifying significant polygenic overlap with numerous psychiatric, neurodevelopmental, and neurodegenerative disorders^11,12,14,15^. However, cortical expansion is not synonymous with surface area, as the latter does not capture the gyrification and folding of growing cortical tissue. Consequently, studies relying on any single morphometric measure will likely overlook key components of a multi-faceted neurodevelopmental process.

A more nuanced approach is therefore needed—one that appropriately models the multiple features that jointly index cortical expansion. Genomic Structural Equation Modeling^16^ (Genomic SEM) has emerged as a rigorous statistical framework for modeling the shared genetic architecture of related traits. Using this approach, GWAS results for several T1-weighted imaging-derived phenotypes were modeled as a latent genomic factor underlying cortical expansion. Warrier and colleagues previously demonstrated that this cortical expansion factor could be estimated from global indices of folding, curvature, gyrification, surface area, and volume, and it was genetically distinct from other macro- and microstructural dimensions of cortical morphometry^13^. However, the specific loci and genes associated with this cortical expansion factor, its regional heterogeneity across the cortex, and its relationships with neuropsychiatric disorders remain unknown.

Here, we build upon recent work and model a global cortical expansion factor using GWAS data from more than 70,000 individuals, mapping its genetic architecture across multiple levels of analysis and its polygenic relationships with diverse neuropsychiatric conditions. We also examine genetic influences at a finer-grained scale, characterizing the genetic topography of expansion across distinct cortical regions to identify functionally relevant patterns of heterogeneity and pleiotropy. Collectively, our results provide new insight into the developmental neurobiology of cortical expansion and its relationship with human health.

## Results

### Tangential growth and gyrification are influenced by a shared pleiotropic architecture

To characterize common variant genetic influences on cortical expansion, we obtained GWAS results for five global MRI-derived phenotypes: folding index (FI), intrinsic curvature index (ICI), local gyrification index (LGI), surface area (SA), and intracranial volume (ICV) (**Table 1**; **Methods**). Using LD score regression^17^ and MiXeR^18^, we found that all traits had statistically significant and comparable SNP heritability (0.27 [s.e. = 0.02] to 0.37 [s.e. = 0.03]) and polygenicity estimates (1,744 [s.d. = 105] to 2,224 [s.d. = 153] causal variants), with SA being the most heritable and FI being the most polygenic (**Table S1**). Moreover, we found that all imaging-derived phenotypes were strongly genetically correlated with each other, though ICV was generally less correlated with the other traits (ICV comparisons: *r*_g_ = 0.42–0.85; all other comparisons: *r*_g_ = 0.71–0.93; **Fig. 1a**; **Table S2**; **Fig. S1**). Further inspection revealed that these patterns of polygenic overlap generally reflected a combination of strong concordant pleiotropy, modest discordant pleiotropy, and limited trait-specific architecture (**Fig. 1b**).

**Figure 1.**
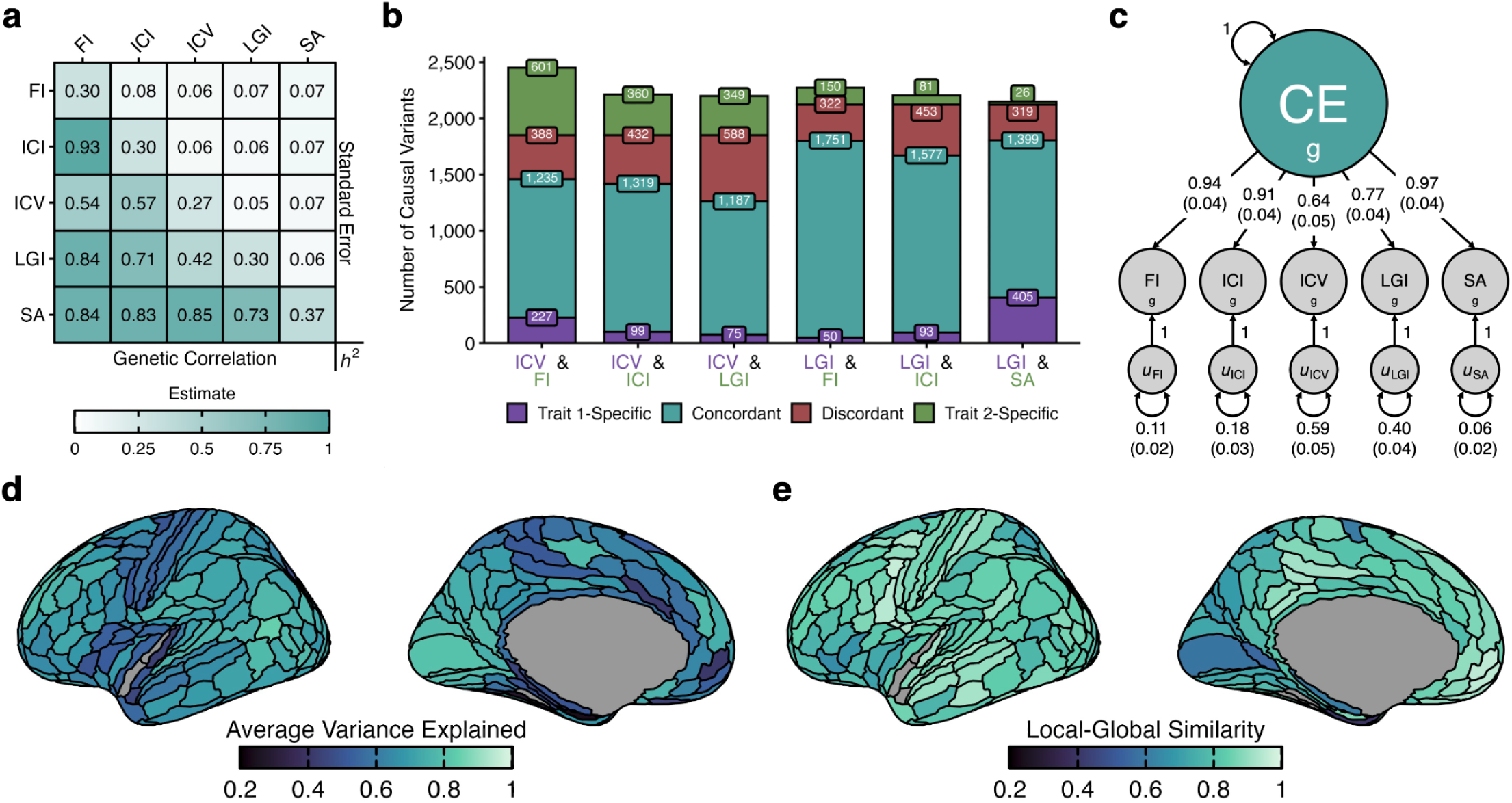
Multivariate genetic architecture of cortical expansion. **a)** Genetic correlation matrix between folding index, intrinsic curvature index, local gyrification index, surface area, and intracranial volume. **b)** Overlapping polygenic architecture between imaging traits using bivariate MiXeR. The scale represents the estimated number of causal variants per trait at 90% SNP heritability. Trait 1 / 2-Specific: Variants influencing only one trait (Trait 1 corresponds to the first trait on the x-axis label). Concordant: Variants affecting both traits in the same direction. Discordant: Variants affecting both traits in opposite directions. **c)** Structural equation model path diagram for a global cortical expansion (CE) model. **d)** Average indicator variance explained (AVE) by regional cortical expansion factors, as measured by the mean of the squared loadings (variance explained) across indicators for each region (**Methods**). **e)** Genetic correlation (r_g_) between regional cortical expansion factors and the global cortical expansion factor. Gray regions in **d)** and **e)** reflect those that were excluded due to negative heritabilities in ICI and FI data (**Methods**).

**Table 1.** Summary of GWAS summary statistics used in global cortical expansion factor model. The sample size (*N*) reflects the maximum of the SNP-level sample sizes for each imaging-derived phenotype. The heritability (*h*^2^), genomic inflation factor (*λ*_GC_), statistical power, LD Score regression intercept, and LD Score regression ratio were estimated using LD Score regression in Genomic SEM. *λ*_GC_ represents the median *χ*^2^ statistic divided by the expected median of the *χ*^2^ with one degree of freedom. Mean *χ*^2^ refers to the average *χ*^2^ statistic across SNPs and represents statistical power. The intercept is reported from the LD Score regression intercept, while the ratio refers to 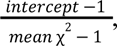 which can be interpreted as a measure of bias attributable to population stratification. For all LD Score regression analyses, we used European-like HapMap3 SNPs with MAF ≥ 0.01. Polygenicity was estimated as the number of causal variants per component at 90% SNP heritability using MiXeR. Errors for polygenicity represent the standard deviations across twenty random iterations of MiXeR, using a random set of 600,000 SNPs each iteration.

| GWAS (abbr.) | Source | *N* | $h^2$ | $\lambda_{GC}$ | Mean $\chi^2$ | Intercept | Ratio | Polygenicity |
| --- | --- | --- | --- | --- | --- | --- | --- | --- |
| Surface area (SA) | Warrier et al., 2023<sup>13</sup> | 36,663 | 0.37 | 1.20 | 1.29 | 1.01 | 0.05 | 1744.28 (105.29) |
| Folding index (FI) | Warrier et al., 2023<sup>13</sup> | 36,663 | 0.30 | 1.17 | 1.24 | 1.01 | 0.04 | 2223.73 (153.45) |
| Intrinsic curvature index (ICI) | Warrier et al., 2023<sup>13</sup> | 36,663 | 0.30 | 1.16 | 1.23 | 1.01 | 0.05 | 2111.27 (116.90) |
| Local gyrification index (LGI) | Warrier et al., 2023<sup>13</sup> | 36,663 | 0.30 | 1.17 | 1.22 | 1.00 | 0.01 | 2123.35 (134.53) |
| Intracranial volume (ICV) | Garcia-Marin et al., 2024<sup>24</sup> | 73,800 | 0.27 | 1.26 | 1.42 | 1.01 | 0.03 | 1849.55 (79.28) |

Using Genomic SEM, we proceeded to test our hypothesis that the observed patterns of pairwise genetic overlap could be explained by a polygenic architecture that jointly influences all five morphometric traits. We found evidence of such an architecture: a common factor model showed good fit to the data (*χ*^2^(5) = 471.37; Akaike information criterion (AIC) = 491.37; comparative fit index (CFI) = 0.96; standardized root mean square residual (SRMR) = 0.08), with large standardized loadings for all indicators (λ = 0.64–0.97; **Fig. 1c**; **Table S3**).

Given our goal of region-specific modeling, we also evaluated a cortical expansion factor that omitted ICV, which has no regional counterpart (**Methods**). This adjusted global cortical expansion model showed good fit to the data (*χ*^2^(2) = 100.65; AIC = 116.65; CFI = 0.99, SRMR = 0.02; **Fig. S2**) with similar factor loadings (λ = 0.82–0.99; maximum Δλ = 0.09). When examining patterns of genetic overlap at the level of individual cortical regions based on the Human Connectome Project parcellation^13,19^, we found that a common factor model exhibited acceptable fit (CFI > 0.9; SRMR < 0.1) in ∼88% (155/177) of tested regions (**Fig. S3**). Inspection of the residual covariance matrix revealed that model misfit in 21 regions not achieving acceptable fit was driven by stronger genetic coupling between ICI and FI (16/21) or between SA and LGI (5/21). After allowing for these pairs to covary in a region-specific manner (**Methods**), acceptable model fit was observed across the cortex (**Fig. S3**). Across all regions, the cortical expansion factors explained a substantial portion of the indicator variance (median 67.65%; range 23.59-84.67%) (**Fig. 1d**), as measured by the mean of the squared indicator loadings (**Methods**). Regional factor loadings were broadly similar to those observed in the global model (**Fig. S4**; **Table S4**). However, regional LGI measures were generally more modestly related to cortical expansion (mean *λ* = 0.52) than the global LGI measure (*λ* = 0.82), particularly in the parietal cortex, suggesting that some patterns of sulcal complexity are influenced by more region-specific genetic architectures.

To further interrogate regional genetic heterogeneity in cortical expansion, we estimated the genetic correlation between each of the regional factors and the global factor. All regions were at least moderately genetically correlated with the global factor (*r*_g_ = 0.40–0.97; **Fig. 1e**). We then characterized the topographical organization of local-global genetic similarity by computing the spatial correlation between the local-global map and 10 canonical cortical maps indexing anatomical, functional, developmental, and evolutionary gradients (**Methods**)^20,21^. We tested significance using spin tests^22,23^ and found a significant spatial correlation between the local-global similarity map and the developmental expansion map (Spearman’s *ϱ* = 0.43, *P* = 0.002; **Fig. S5**), indicating that regions with the strongest local-global coupling also tended to be those that undergo the most profound expansion during development. Overall, these results indicate that the polygenic and pleiotropic architecture underlying human cortical expansion can be parsimoniously modeled as a common genetic factor at both the global and regional level.

### Genes subserving cortical expansion implicate neurodevelopmental and neurodegenerative processes

To identify the specific genetic variants associated with cortical expansion, we conducted a multivariate GWAS of the global cortical expansion factor, aggregating information across a combined 220,452 observations from at least 73,800 unique individuals (**Fig. 2a**) (**Methods**). We observed substantial inflation of the association statistics (*λ*_GC_ = 1.26, mean *χ*^2^ = 1.35) that was primarily attributable to polygenic signal rather than population stratification or related confounding (LD Score intercept = 1.02 [s.e. = 0.01], attenuation ratio = 0.06 [s.e. = 0.03]) (**Fig. S6**). As a sensitivity analysis, we repeated the multivariate GWAS after excluding ICV and observed a highly similar pattern of results (**Table S5**); nonetheless, we retained ICV in our primary model as its inclusion improved statistical power for discovery (**Methods**).

**Figure 2.**
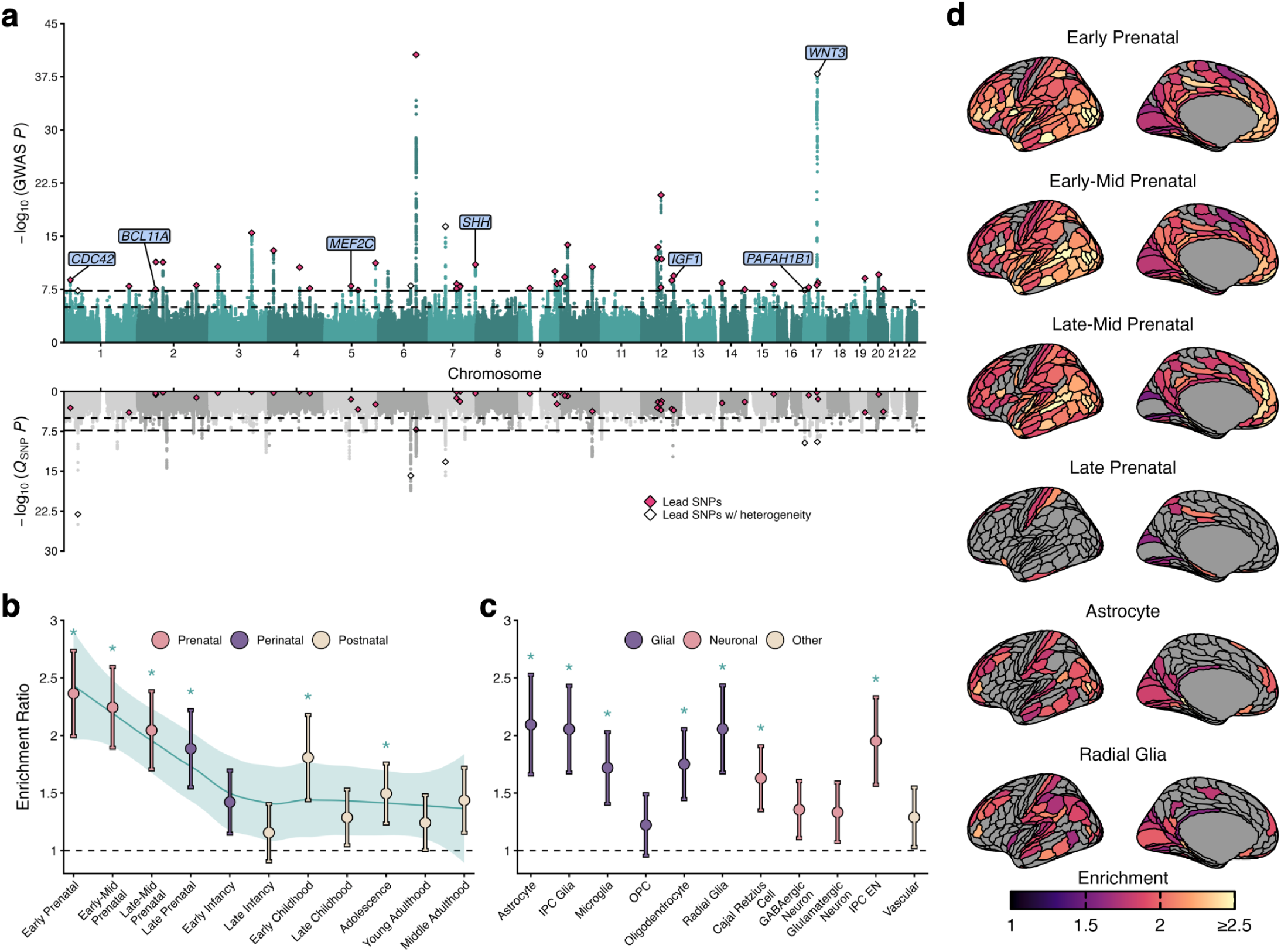
Identification of genetic variants and enrichment of global cortical expansion. **a)** Multivariate GWAS for the cortical expansion factor. Diamonds represent lead SNPs for loci identified by FUMA. Annotated genes include seven effector genes (cumulative precision ≥ 0.75) that were identified via FLAMES and overlap with the Deciphering Developmental Disorders Study. **b)** Enrichment for developmental annotations in Stratified Genomic SEM, where error bars reflect the 95% confidence interval of the enrichment estimate. **c)** Enrichment for cellular annotations in the neocortex during the first trimester in Stratified Genomic SEM, where error bars reflect the 95% confidence interval of the enrichment estimate. **d)** Regional enrichment for select annotations from developmental epochs and neocortical cells, where only regions with Bonferroni-corrected significant enrichment are colored. IPC Glia: intermediate progenitor cell for glia; IPC EN: intermediate progenitor cell for glutamatergic excitatory neurons; OPC: oligodendrocyte precursor cell.

We subsequently used FUMA^25^ to characterize the variant-level signals associated with cortical expansion, identifying a total of 48 approximately independently linked genomic loci with 56 lead SNPs (**Table S6**). The most significant loci included rs8756 (*Z* = 9.53, *P* = 1.56e-21), a 3′ UTR SNP in *HMGA2*, which plays a role in fetal and postnatal growth^26^; and rs1378358 (*Z* = 13.00, *P* = 1.27e-38), an intronic SNP located within the 17q21.31 cytogenetic region, which harbors a 900 kb inversion region that has previously been associated with individual variation in cortical morphology and neurodegenerative diseases^27^. Six of the lead SNPs (rs11629584, *Z* = -5.81, *P* = 6.09e-9; rs11652522, *Z* = 5.77, *P* = 8.16e-9; rs11978532, *Z* = -5.72, *P* = 1.04e-8; rs2106164, *Z* = -5.84, *P* = 5.25e-9; rs4672381, *Z* = 5.54, *P* = 3.08e-8; rs7025791, *Z* = -5.87, *P* = 4.31e-9) did not reach genome-wide significance in any of the univariate summary statistics comprising the cortical expansion factor. Notably, most associated lead SNPs (51/56) had non-significant *Q*_SNP_ test statistics (**Fig. 2a**; **Fig. S6**) (**Methods**), including all six of the novel lead SNPs, indicating that the common pathway represents a plausible model of pleiotropy. Moreover, 95% (53/56) of associated lead SNPs remained significant in sensitivity analyses that adjusted for heterogeneity identified via *Q*_SNP_ (**Fig. S7**) (**Methods**). We further applied fine-mapping via SuSiE^28^ and identified 54 unique 95% credible sets with a purity of at least 0.5 across 45 genomic loci (**Table S7**).

We then used MAGMA^29^ to collate SNP-level signals into gene-level signals to aid in the identification of genes and biological pathways associated with cortical expansion. Briefly, we found that 122 of the 17,276 tested genes were significant at the genome-wide threshold. Subsequent MAGMA-based gene-set enrichment analysis revealed that the polygenic signal associated with cortical expansion was enriched in seven pathways with established ties to regulatory and developmental processes (**Table S10**), including those involved in Wnt signaling, which is an evolutionarily conserved pathway responsible for coordinating crucial cell functions and development^30^, and the cell cycle^31^.

Finally, we used FLAMES, which combines SNP-to-gene evidence and convergence-based evidence, to predict putative effector genes that mediate locus-trait associations^32^. We identified 30 unique effector genes across 35 genomic regions with cumulative precision ≥ 0.75, as well as one high-confidence effector gene (*MEF2C*; cumulative precision ≥ 0.999) (**Table S8**). Notably, we found that effector genes were enriched in genes definitively implicated in developmental disorders from the Deciphering Developmental Disorders Study^33,34^ (*P* = 0.002) (**Table S9**) (**Methods**). Overlapping effector genes included those responsible for intellectual disability (*BCL11A*, *CDC42*, *MEF2C*) and cerebral malformations (*MEF2C*, *PAFAH1B1*, *SHH*). Consistent with our MAGMA results, *WNT3*, the deletion of which causes WNT3-related tetra-amelia syndrome, was also involved, further implicating the Wnt signaling pathway in cortical expansion. Notably, *PAFAH1B1* captures neurobiology specific to cortical expansion, as its absence causes *PAFAH1B1*-related lissencephaly, a severe malformation of neurodevelopment characterized by absent or markedly reduced cortical folding.

### Broadly distributed genetic architecture is evolutionarily conserved and developmentally sensitive

To better understand the forms of genomic variation that subserve cortical expansion, we used Stratified Genomic SEM^35^ to test for enrichment across 69 total annotations, including 46 baseline annotations^36,37^, 11 developmental annotations from BrainSpan^38^, and 12 annotations of neocortical cell-type-specific gene expression in the first trimester^39,40^ (**Methods**).

Briefly, genes with a high probability of loss-of-function intolerance were enriched (Enrichment Ratio = 2.16 [s.e. = 0.14]; *P* = 1.63e-16), indicating the involvement of core biological processes^41^. We also observed significant enrichment in nine baseline annotations for the global cortical expansion factor, including those that harbor human-specific, ancient, and evolutionarily conserved sequences (*P* < 0.05 / 69 = 7.25e-4, **Fig. S8**)^42^. In addition, we found significant enrichment in several regulatory genomic annotations, including various markers of active gene regulation^43^ and super-enhancers^44^. This pattern is consistent with the idea that cross-species, conserved neurodevelopmental genes are complemented by human-specific changes in regulatory elements to drive evolutionary changes^45^. With respect to the developmental annotations, the polygenic signal for global cortical expansion was most prominently enriched in genes highly expressed during the earliest developmental epochs—spanning early to late prenatal development—but also in those that were highly expressed during early childhood and adolescence (**Fig. 2b**, **Fig. S9**). Notably, when testing for enrichment in human prenatal neocortical cell types, we found robust evidence of enrichment across the three major types of glial cells (astrocytes, oligodendrocytes, microglia), Cajal-Retzius cells, radial glial cells, and broader progenitor cell types (**Fig. 2c**). Significant enrichment of radial glial cells and other progenitor cells bolsters our MAGMA- and FLAMES-based findings involving Wnt signaling, as the Wnt signaling pathway regulates the proliferation and differentiation of these cells^46^. Additional sensitivity analyses evaluated the impact of annotation size (**Fig. S10**) and annotation formation (**Fig. S11**) (**Methods**) and found similarly strong enrichment in progenitor cell types.

Given the regional heterogeneity of genetic influences on cortical expansion demonstrated above, we used Stratified Genomic SEM to evaluate whether enrichment signals exhibited meaningful patterns of interregional variation. After correcting for multiple comparisons across regions, we observed widespread significant enrichment in the early-, early-mid- and late-mid-prenatal annotations, underscoring the importance of developmentally sensitive genes in expansion across the cortex **(Fig. 2d**; **Fig. S12a**). Interestingly, enrichment of the cortical expansion signal in genes expressed in the late prenatal period was more selective, suggesting that perinatal neurodevelopmental biology may be distinct from earlier prenatal periods. With respect to the neocortical cell-type annotations, we found that enrichment patterns were most widespread in glial cell types and progenitor cells (**Fig. S12b**). In particular, widespread enrichment of radial glia and astrocytes reflects their prominent roles in the gyrification and folding of the cortex^47–49^ (**Fig. 2d**).

To delineate the topographical organization of regional enrichment patterns, we used spin tests^22,23^ to characterize the spatial correlation between each pair of annotations and compared to canonical maps (**Methods**). Here, we found that spatial topography of enrichment patterns for the early-, early-mid-, and late-mid developmental annotations were highly similar (**Fig. S13**). We also found that the pattern observed for genes highly expressed during the late prenatal period was distinct from earlier periods, instead being more similar to the early infancy annotation, further suggesting potentially distinct biology of the perinatal period. Among neocortical cell annotations, genes expressed in radial glia demonstrated similar spatial patterning to astrocytes and intermediate progenitor cells for glia and excitatory neurons (**Fig. S14**). To contextualize these findings along anatomical, functional, developmental, or evolutionary axes, we performed spin tests with 10 canonical maps (**Fig. S12c**; **Fig. S5b** for canonical maps). Among both developmental and cell-type annotations, enrichment tended to be in areas that were most genetically coupled with the global signal (**Fig. 1e**). Enrichment in genes during late infancy followed the sensorimotor-association axis, potentially reflecting a critical developmental window during which association regions of the brain differentially expand (**Fig. S12c**). Collectively, these enrichment patterns implicate early development, including broad neurobiological pathways underlying prenatal neurodevelopment and diverse progenitor cell lineages.

### Genetic associations with cortical expansion are shared across diverse brain-based disorders

We next used Genomic SEM to characterize the degree of genetic sharing between cortical expansion and 20 neuropsychiatric disorders (**Table S13**). Briefly, we found that the global cortical expansion factor was negatively genetically correlated with attention-deficit/hyperactivity disorder (ADHD), tobacco use disorder (TUD), posttraumatic stress disorder (PTSD), cannabis use disorder (CUD), and neuroticism, and positively genetically correlated with Parkinson’s disease (**Fig. 3a**). *Q_Trait_* analyses suggested that these genomic relationships plausibly operated through the model-implied common pathway **(Fig. S15**) (**Methods**).

**Figure 3.**
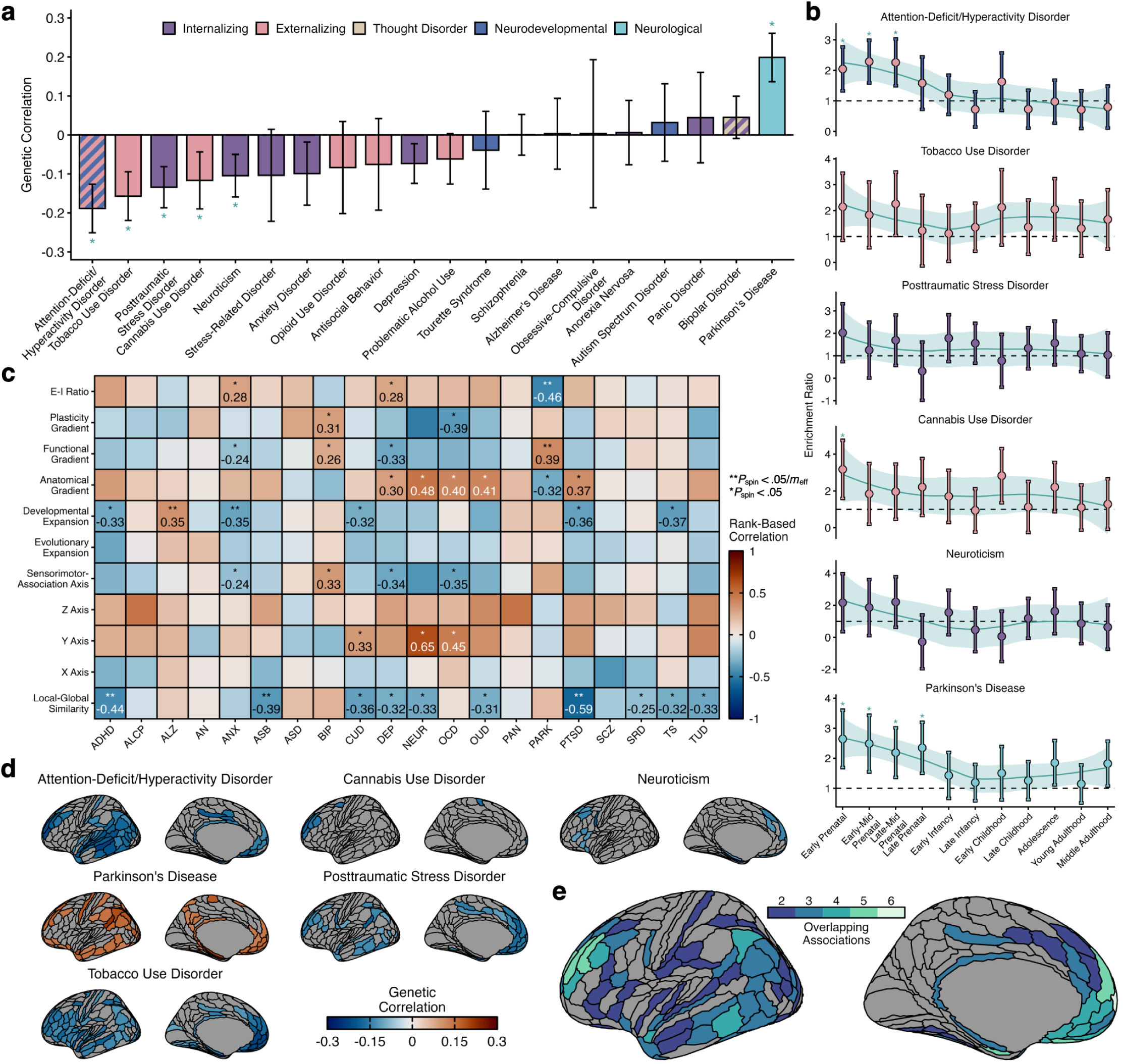
Relationship between cortical expansion and neuropsychiatric disorders. **a)** Genetic correlations between the cortical expansion factor and various psychiatric traits/disorders, where error bars reflect the 95% confidence interval of the genetic correlations. Due to data access stipulations, ICV was excluded from any analysis of ALCP, ASB, CUD, OUD, or TUD; a comparable four-indicator cortical expansion factor model was used instead in those cases. **b)** Enrichment of the covariance between cortical expansion and ADHD, TUD, PTSD, CUD, NEUR, and Parkinson’s in genes expressed in eleven developmental epochs, where error bars reflect the 95% confidence interval of the enrichment estimate. An asterisk (*) reflects significant enrichment after Bonferroni correction across 11 developmental annotations. (**c)** Comparison of unthresholded regional correlation maps to canonical maps. A single asterisk (*) reflects *P_spin_* < 0.05, and a double asterisk (**) reflects significance corrected for the effective number of tests (see descriptions in **Methods**). **d)** Regional correlation maps between cortical expansion and ADHD, TUD, PTSD, CUD, NEUR, and Parkinson’s, thresholded by correcting for the number of effective tests across regions. **e)** Consensus maps depicting the number of disorders for which each region was significant, derived from **d)**. ADHD: attention-deficit/hyperactivity disorder; ALCP: problematic alcohol use; ALZ: Alzheimer’s disease; AN: anorexia nervosa; ANX: anxiety disorder; ASB: antisocial behaviors; ASD: autism spectrum disorder; BIP: bipolar disorder; CUD: cannabis use disorder; DEP: depression; NEUR: neuroticism; OCD: obsessive-compulsive disorder; OUD: opioid use disorder; PAN: panic disorder; PARK: Parkinson’s disease; PTSD: post-traumatic stress disorder; SCZ: schizophrenia; SRD: stress-related disorders; TS: Tourette’s syndrome; TUD: tobacco use disorder.

Using Stratified Genomic SEM, we tested whether the genetic covariance between cortical expansion and these six disorders was enriched in specific annotations. Here, we found that the covariances between cortical expansion and ADHD, CUD, and Parkinson’s were all enriched for genes highly expressed during the prenatal epoch **(Fig. 3b**). In addition, we found that several of the developmental enrichment patterns were similar across disorders. Three externalizing spectrum disorders (ADHD, CUD, and TUD) demonstrated highly correlated patterns of enrichment (*r* = 0.60–0.71) in their covariance with cortical expansion, with the strongest enrichment observed in prenatal annotations. Interestingly, the developmental enrichment trend for Parkinson’s was also similar to those three externalizing disorders (*r* = 0.50–0.79). Among genes expressed in neocortical cell types, the covariance between cortical expansion and CUD was enriched in radial glia, while the TUD covariance was enriched in oligodendrocytes (**Fig. S18**).

Given the observed regional heterogeneity of genetic influences, we also considered the possibility of regional differences in the relationship between cortical expansion and neuropsychiatric disorders (**Fig. S16**; **Fig. S17**). After generating cortex-wide genetic correlation maps for each of the 20 disorders (**Methods**), we found that the strongest genetic correlations tended to be concentrated in regions that (i) undergo greater expansion across development, (ii) exhibit lower T1w/T2w ratio (a proxy measure of reduced myelination) along the anatomical gradient, (iii) show decreases in excitatory-inhibitory balance during adolescence^21^, and (iv) are preferentially situated in association regions of the sensorimotor-association axis (**Fig. 3c**). The six phenotypes significantly genetically correlated with global cortical expansion also exhibited the greatest number of significant regional correlations, ranging from seven (CUD) to 88 (TUD) significant regions (**Fig. 3d**). When evaluating overlap across these brain-disorder associations, we found the greatest cross-disorder significance in the prefrontal cortex (**Fig. 3e**), a region that has been implicated across multiple neuropsychiatric disorders^50^.

### Genetic overlap with clinical conditions reflects a nuanced mixture of effects

Since a mixture of concordant and discordant pleiotropy may result in attenuated or even near-zero genome-wide genetic correlations, we also used the MiXeR^18,51^ framework to interrogate the polygenic overlap of cortical expansion and the 20 neuropsychiatric disorders. In univariate MiXeR analyses, we observed good model fit for all but six phenotypes (antisocial behavior, anxiety disorder, panic disorder, obsessive-compulsive disorder, Tourette’s syndrome, and stress disorder), for which the causal mixture model provided worse fit (larger AIC and Bayesian information criterion (BIC)) than the infinitesimal model (**Table S11**). These analyses yielded polygenicity estimates for all tested phenotypes. We observed the highest polygenicity estimates for depression, followed by other psychiatric disorders, and lowest polygenicity estimates for neurological disorders (Alzheimer’s disease, Parkinson’s disease). Notably, among well-fitting models, cortical expansion had lower polygenicity than all psychiatric and developmental disorders, but higher polygenicity than both neurological disorders (**Fig. S19**), consistent with recent findings regarding the polygenicity of surface area^52^.

We subsequently carried forward phenotypes with valid univariate MiXeR results into bivariate MiXeR analyses, which identified high degrees of overlap between cortical expansion and most tested phenotypes (**Fig. 4**). Model fits were largely stable across random iterations (**Fig. S20**). The models all demonstrated better fit than the minimally overlapping model. However, eight models (neuroticism, cannabis use disorder, tobacco use disorder, post-traumatic stress disorder, schizophrenia, autism spectrum disorder, anorexia nervosa, opioid use disorder) had greater AIC (worse fit) than the maximally overlapping model (**Table S12**). These cases reflect convergence to maximal polygenic overlap, suggesting a high degree of shared polygenic architecture. Due to the differing polygenicities of neurological disorders, psychiatric disorders, and cortical expansion, we estimated the extent of overlap as the ratio of shared variants to the number of causal variants for the less polygenic trait of each pair (**Methods**). This median proportion of shared variants for the less polygenic trait of each pair was 82.24% (range: 50.01–97.03%). The modest or absent genome-wide correlations mask highly overlapping genetic architecture between cortical expansion and neuropsychiatric disorders that is driven by a mixture of concordant and discordant effects, revealing transdiagnostic, pleiotropic SNPs that may share cortical expansion as a common etiological axis.

**Figure 4.**
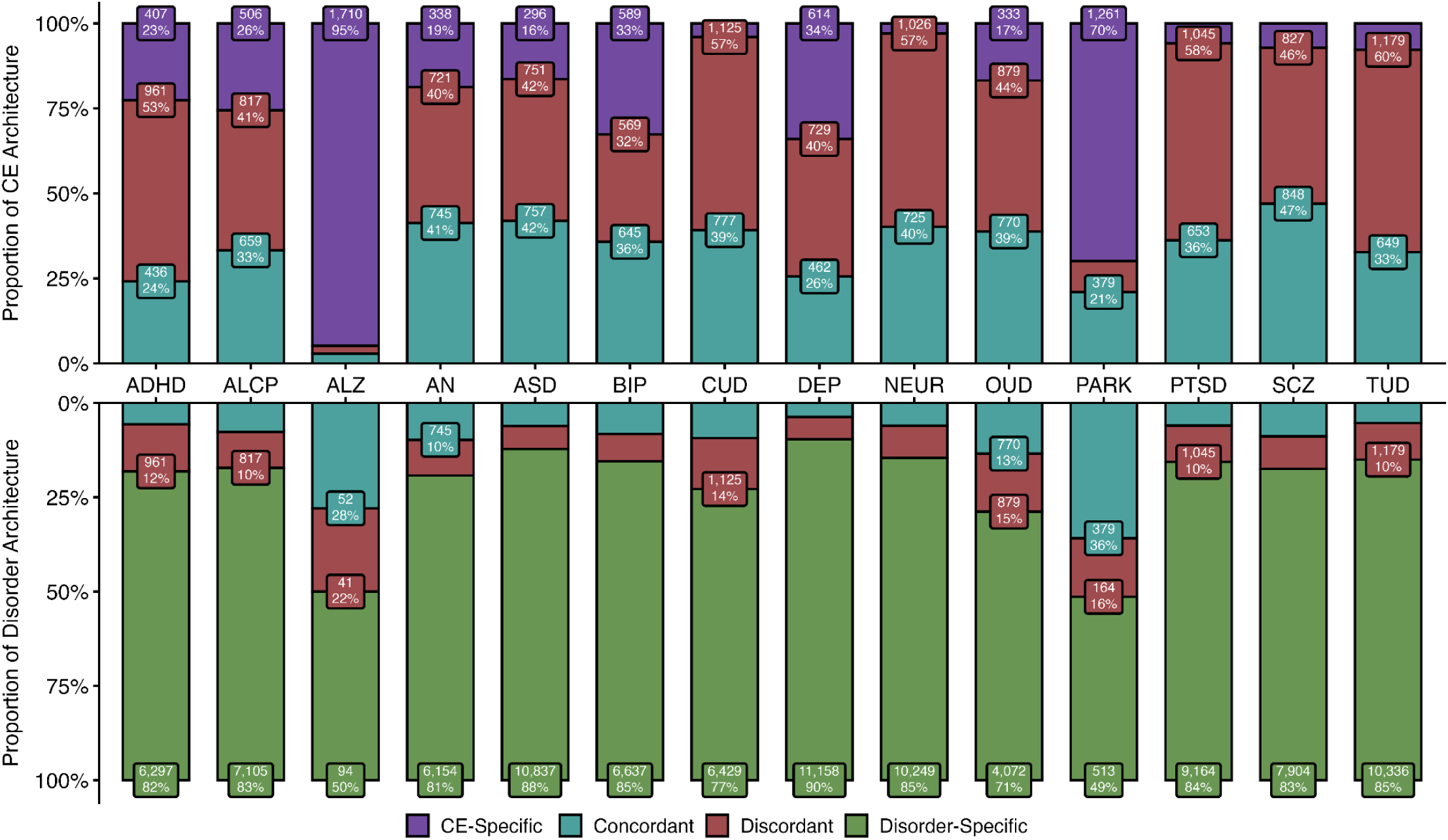
Shared polygenic architecture via MiXeR. Bivariate MiXeR results show causal genetic overlap between cortical expansion (CE) and disorders. The top panel reflects the proportion of CE polygenic architecture partitioned by CE-specific variants and variants shared with each neuropsychiatric disorder, further partitioned by concordance or discordance. Similarly, the bottom panel reflects the proportion of neuropsychiatric polygenic architecture partitioned by disorder-specific variants and variants shared with CE, further partitioned by concordance or discordance. Numbers on each component represent the estimated number of causal variants per component at 90% SNP heritability. Note that, for the purposes of plotting, component values are only reported for components ≥ 10%. Due to data access stipulations, ICV was excluded from any analysis of ALCP, ASB, CUD, OUD, or TUD; a comparable four-indicator CE factor model was used instead in those cases. ADHD: attention-deficit/hyperactivity disorder; ALCP: problematic alcohol use; ALZ: Alzheimer’s disease; AN: anorexia nervosa; ASD: autism spectrum disorder; BIP: bipolar disorder; CUD: cannabis use disorder; DEP: depression; NEUR: neuroticism; OUD: opioid use disorder; PARK: Parkinson’s disease; PTSD: post-traumatic stress disorder; SCZ: schizophrenia; TUD: tobacco use disorder.

## Discussion

Cortical expansion is a fundamental neurodevelopmental program that has been central to human evolution, shaping a key structural substrate for higher-order cognition. Yet, despite convergent evidence linking variation in cortical morphology to human health and disease, the shared genetic architecture underlying the tangential growth and gyrification of the cortex has remained poorly understood. Here, we address this gap by characterizing common variant influences on cortical expansion, charting the developmental biology of this process through a genomic lens and mapping its genetic relationships with a diverse set of neuropsychiatric disorders. Taken together, our findings support several novel insights into the genomic architecture of cortical expansion, its heterogeneity across the cortex and development, and its relevance to shared liability across brain-based disorders.

Building upon recent work^13^, we show that a latent genomic factor underlying global cortical expansion explains genetic sharing among five morphometric features: folding index, intrinsic curvature index, local gyrification index, surface area, and intracranial volume. Although these features measure methodologically and conceptually distinct aspects of neuroanatomical variation, our results reveal that they are largely shaped by a shared genetic architecture. For the first time, we extend this framework to the regional level, demonstrating that analogous local expansion factors account for much of the genetic overlap among folding, curvature, gyrification, and surface area. Patterns of regional genetic sharing were not spatially uniform, though; we observed substantive regional heterogeneity across the cerebral mantle. Regions that undergo the most profound expansion during development tended to be influenced by genetic architectures more similar to that of the whole cortex, but others showed more differentiated local architectures. Our results indicate that regionally specific effects on sulcal complexity were a key contributor to this pattern of genetic differentiation.

We also characterize the functional genomic architecture of cortical expansion across several biological scales. First, using annotation-based genomic partitioning, we found robust enrichment in genes specifically expressed during three broad developmental periods: pre- and perinatal development, early childhood, and adolescence. Notably, these patterns roughly approximate those observed for the phenotypic expansion of cortical surface area^4^, which begins rapidly developing in the prenatal period, reaches peak velocity shortly after birth, and peaks in size around the turn of adolescence. Moreover, we found enrichment in multiple evolutionarily conserved/ancient sequences^36^, which could align with the hypothesis that all mammalian species trace back to a common ancestor with a gyrencephalic cortex^31,53^. Indeed, taken together, these findings suggest the architecture of human cortical expansion may center on deeply conserved neurodevelopmental programs, alongside enhancer- and promoter-linked regulatory elements that may have helped shape cortical organization.

At a finer level of biological resolution, prominent patterns of enrichment were observed in progenitor cell annotations, including radial glia, implicating proliferative and scaffolding processes as central components of cortical expansion^3,31,49,54,55^. This pattern aligns with the radial unit hypothesis, which asserts that neuronal cells migrate from the same place of origin along a common radial glial scaffold and settle in the same radial column^56–58^, although more recent studies have demonstrated complexities beyond strictly radial migration^2^. In this framework, variation in the pool of progenitor cells and radial glial organization can influence the tangential growth and gyrification of the cortex. However, enrichment was not restricted to progenitor cell populations. We also found enrichment in genes specifically expressed in glial and neuronal cell types related to cortical development and organization. Specifically, we observed enrichment in astrocytes, consistent with the role of astrogenesis in gyrification of the cortex^47,48^, as well as oligodendrocytes, microglia, and Cajal-Retzius cells. Thus, rather than localizing to a single privileged cell class, common variant influences on cortical expansion appear to be distributed across a broader, developmentally sensitive cellular ensemble.

Our analyses also afforded new insights into the specific molecular pathways underlying human cortical expansion. Across multiple analyses, Wnt signaling emerged as a recurring theme. For instance, our competitive gene-set analyses implicated three gene sets related to Wnt signaling, and effector gene mapping identified *WNT3* as a putatively causal gene. Paired with the enrichment of multiple progenitor cell types, for which Wnt signaling is a key regulator^59^, our results triangulate on this pathway as a contributor to human cortical expansion. Collectively, these results build upon prior research linking genetic influences on surface area to Wnt signaling^11,12^, suggesting that the enrichment of this pathway is not exclusive to areal growth but also extends to aspects of gyrification and curvature.

Converging lines of evolutionary, clinical, and observational evidence implicate atypical cortical expansion as a transdiagnostic liability for neurocognitive dysfunction, broadly construed^8–10^. Our findings provide additional support for this hypothesis. Specifically, we found that the common variant architecture of cortical expansion exhibited substantial polygenic overlap with diverse neuropsychiatric disorders. Notably, even when no significant genetic correlation was observed, bivariate causal mixture modeling revealed substantial shared genetic architecture. Viewed in the context of recent psychiatric and neurological genetics studies, these findings suggest that cortical expansion may index a neurodevelopmentally anchored substrate through which pleiotropic liability is distributed across brain-based disorders—both within and across clinical domains^52,60^.

While polygenic overlap between cortical expansion and clinical conditions typically involved complex mixtures of concordant and discordant pleiotropy, several disorders exhibited directionally consistent patterns of genetic sharing. Generally, reduced cortical expansion was significantly associated with increased liability for neuropsychiatric disorders, though the trend was reversed for Parkinson’s, corroborating previous studies of the relationship between Parkinson’s and surface area^11,61,62^. Although the mechanism underlying this relationship remains unclear, these findings add to mounting evidence of shared etiology between neurodevelopment and Parkinson’s disease^63,64^. Regardless of the direction of effect, when these genetic correlations were identified, we found that they were distributed in a topographically meaningful manner. That is, cortical regions with the strongest genomic links to clinical phenotypes were relatively conserved across disorders, despite their differing developmental trajectories and clinical presentations. These relationships tended to localize in association regions relative to sensorimotor regions, as well as those with lower myelination, as approximated by T1w/T2w ratio. In this sense, our findings add to recent evidence that the genetics of surface area and psychiatric disorders are related along an axis of anatomical hierarchy^14^, and also add the possibility of certain neurodegenerative conditions, such as Parkinson’s disease, operating along a similar hierarchy.

While our study provides a useful genomic framework for understanding cortical expansion, it is not without limitations. First and foremost, our analyses were restricted to individuals of European-like ancestry, potentially limiting the generalizability of our findings. Although imaging genetics resources are gradually becoming more diverse^65^, data from non-European-like ancestry groups are currently very limited. Second, our study generates developmental biological insight by studying the genomic architecture of imaging-derived phenotypes influenced by cortical expansion rather than directly modeling the developmental process itself. Nevertheless, our findings both recapitulate and extend established developmental neurobiology, bolstering the validity of etiological insights generated with this approach. Third, multiple analyses rely on results from the multivariate GWAS (e.g., FLAMES), which we only performed globally. Although we investigated spatial heterogeneity via Genomic SEM, an exhaustive investigation of spatial heterogeneity of multivariate GWAS remains beyond the objectives of this work. Fourth, our analyses characterizing polygenic overlap with clinical conditions leveraged publicly available GWAS that often relied on heterogeneous case definitions and ascertainment strategies. It is possible that other clinical phenotypes, such as specific symptoms, would yield different patterns of overlap, but that is beyond the scope of this work. Fifth, our custom genomic annotations are based on arbitrary thresholds, such as the commonly employed top decile of expression, although we have reported sensitivity analyses detailing the impact of this analytic decision.

In summary, our findings help elucidate the genomic architecture of cortical expansion and its relevance to a diverse set of brain-based disorders. Rather than relying on any single neuroanatomical proxy, we demonstrate that the etiology of cortical expansion can be modeled in a latent variable framework that jointly considers the genetic architectures of multiple morphometric features. This approach allowed us to characterize its biology across multiple scales, from variants to genes to pathways to genomic partitions. Through this integrative multivariate modeling framework and complementary bioinformatic analyses, we show that cortical expansion is not just a defining feature of human evolution and development, but a plausible etiological axis linking neurodevelopmental, psychiatric, and neurological conditions across the lifespan.

## Methods

### GWAS data acquisition

Univariate GWAS summary statistics for four MRI-derived phenotypes (SA, FI, ICI, and LGI) were obtained from a previous study^13^ of European-like ancestry individuals from the UK Biobank (*N*_max_ = 31,797) and Adolescent Brain Cognitive Development Study (*N*_max_ = 4,866). Specifically, we obtained GWAS results for a global average of each measure, as well as for 180 bilaterally averaged regions based on the Human Connectome Project parcellation^19^, which defines cortical parcels on the basis of multimodal data (architecture, function, connectivity, and topography). Three regions (52, PHA2, PI) were excluded due to negative heritabilities in ICI and FI data, yielding a total of 177 regions suitable for analysis. Further details are available in the original paper^13^. We also obtained univariate GWAS summary statistics for ICV from a meta-analysis of the ENIGMA consortium, CHARGE consortium, UK Biobank, and ABCD study^24^. In total, there were 73,800 individuals of European-like ancestry. Assuming full sample overlap between the five imaging-derived phenotypes, we conservatively estimated that the combined data included a total of 220,452 observations from at least 73,800 individuals.

Univariate GWAS were obtained for 20 additional external traits, including developmental, psychiatric, and neurological disorders (**Table S13**). The external traits included Attention-deficit/hyperactivity disorder (ADHD)^66^, problematic alcohol use (ALCP)^67^, Alzheimer’s disease (ALZ)^68^, anorexia nervosa (AN)^69^, anxiety disorder (ANX)^70^, antisocial behaviors (ASB)^71^, autism spectrum disorder (ASD)^72^, bipolar disorder (BIP)^73^, cannabis use disorder (CUD)^74^, depression (DEP)^75^, neuroticism (NEUR)^76^, obsessive-compulsive disorder (OCD)^77^, opioid use disorder (OUD)^78^, panic disorder (PAN)^79^, Parkinson’s disease (PARK)^80^, post-traumatic stress disorder (PTSD)^81^, schizophrenia (SCZ)^82^, stress-related disorders (SRD)^83^, Tourette’s syndrome (TS)^84^, and tobacco use disorder (TUD)^85^. Further descriptions of the traits are available in **Table S13.** Note: the ICV summary statistics were excluded from the cortical expansion model for any analysis of ALCP, ASB, CUD, OUD, and TUD due to data access stipulations. Instead, a comparable four-indicator cortical expansion factor model was used in those analyses.

Where necessary, GWASLab was used to liftover GrCh38 GWAS to GrCh37 coordinates, assign rsIDs based on chromosome and base pair number, and identify chromosomes and base pair number for rsIDs^86^. For continuous GWAS, the sample size was considered as the total number of individuals analyzed. For case-control GWAS, we estimated the sum of effective sample size 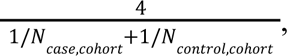 summing across cohorts. If cohort-level case-control data was unavailable, we estimated SNP-level effective sample size as 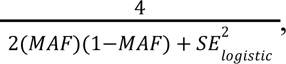 where MAF was either available or estimated using the 1000 Genomes Phase 3 European reference panel^87^. Furthermore, the estimated SNP-level effective sample sizes were bounded between 0.5 and 1.1 times the total effective sample size. These recommendations follow best practices established in prior research^88,89^.

### Polygenicity and discoverability analysis

MiXeR is a statistical framework for characterizing the polygenicity and discoverability of complex traits using GWAS summary statistics. Specifically, univariate MiXeR estimates the polygenicity and discoverability through causal mixture modeling of null and causal components^18^, while bivariate MiXeR is an extension that estimates polygenic overlap by modeling shared causal components, indexing concordant and discordant effects, as well as trait-specific and null components^51^. In this study, MiXeR v1.3 was used to assess polygenic overlap between (i) each pair of univariate imaging summary statistics, and (ii) multivariate cortical expansion with external summary statistics^15,18,51^. The 1000 Genomes Phase 3 data were used as an LD reference panel. As recommended by the developers, we provided the full set of summary statistics for each trait. MiXeR was repeated for 20 random iterations, with 600,000 SNPs randomly selected per iteration. The number of causal variants was based on the proportion explaining 90% of SNP heritability to avoid infinitesimal effects^51^.

The quality of univariate MiXeR models was assessed with AIC relative to the infinitesimal model, calculated by subtracting the AIC of the causal mixture model from the AIC of the infinitesimal model. Any models with negative AIC differences were omitted from follow-up bivariate MiXeR analysis. There were no imaging traits with negative AIC differences (**Table S2**). The quality of bivariate MiXeR models was assessed with the AIC of the minimally overlapping model minus that of the causal mixture model, as well as the AIC of the maximally overlapping model minus that of the causal mixture model. AIC relative to the maximally overlapping model was allowed to be negative as long as the fits were stable across random iterations, which would suggest convergence toward maximal polygenic overlap between two traits^15^. For the imaging traits, we excluded four trait pairs (FI/ICI, ICI/SA, FI/SA, ICV/SA) due to poor fit stability (**Fig. S1**). BIC was also reported but is often overly conservative, so polygenic overlap was considered in spite of a negative BIC^15^.

For bivariate MiXeR models, we calculated the median proportion of shared variants for the less polygenic trait of each pair to assess genetic overlap^52^. For each pair of traits, this metric is calculated as the number of overlapping variants divided by the number of variants for the less polygenic of the two traits in the pair. This value represents estimated overlap relative to the maximal possible overlap, bounded by the less polygenic trait.

### Genomic structural equation modeling

Genomic SEM is a statistical framework for applying structural equation modeling techniques to GWAS summary statistics, facilitating the study of pleiotropic architecture across multiple traits^16^. Briefly, it extends LD Score regression^17^ to the multivariate context by estimating the genetic covariance structure among phenotypes, which can then be formally modeled. Here, we use this approach to model a latent genomic factor indexing cortical expansion and interrogate its architecture via multivariate GWAS and downstream analysis.

Prior to any genomic factor analysis, we first processed data for five imaging-derived phenotypes (FI, ICI, LGI, SA, and ICV) using the *munge* function. Following recommendations of the developers, we restricted these analyses to HapMap3^90^ variants with imputation INFO scores > 0.9 and minor allele frequency > 0.01. We then fit a common factor model to these five imaging-derived phenotypes using Genomic SEM^16^. Parameters were estimated with a diagonally weighted least squares estimator, using unit variance identification to scale the factor. Model fit was assessed using several metrics: the model *χ*^2^ statistic, AIC, CFI, and SRMR. CFI and SRMR were considered as absolute measures of fit^16^; thresholds for acceptable fit were CFI > 0.9^16,91^ and SRMR < 0.1^16,92^, where higher CFI and lower SRMR indicate better fit.

We present this model as an update to previously published work that included GMV as an indicator of cortical expansion^13^, replacing GMV with ICV (*N* = 73,800), which is a conceptually related proxy that is highly correlated with GMV^93^. This change has three distinct statistical and interpretative advantages. First, GMV is approximately the product of SA and cortical thickness (CT). CT is a widely studied imaging-derived phenotype that may be used in future studies in conjunction with cortical expansion, which could cause collinearity issues if GMV was part of the cortical expansion model. Second, the inclusion of ICV brings the number of unique individuals contributing to the cortical expansion factor from 36,663 to 73,800, which is likely to improve statistical power for gene discovery in the Genomic SEM framework. Third, the skull is affected by the expansion of the cortex^2,31^, suggesting it is a relevant proxy measure, and including more proximal measures can improve the psychometric validity of a latent factor^94^. We tested whether including ICV changed the polygenic architecture of cortical expansion by evaluating two common factor models: one including ICV and another excluding ICV.

We also evaluated whether a cortical expansion factor explained patterns of pleiotropy for 177 bilaterally averaged regions. Since ICV is not a regional measure, the region-level common factor only included SA, FI, ICI, and LGI as indicators. While the HCP atlas has 180 regions, three regions (52, PHA2, PI) were excluded due to negative or non-significant heritability for FI and ICI, leaving 177 regions. For an additional three regions (FOP3, s32, Pol1), heritability for FI was not significant, so this term was excluded from the model. Without FI, the model was saturated and thus interpreting model fit was not meaningful, leaving 174 regions to interpret model fit.

For 21 regions not achieving acceptable fit, the models were refit accounting for residual covariance between the two indicators with the highest residual covariance. The highest residual covariance was between ICI and FI for 16 regions, and SA and LGI for five regions. After accounting for this additional regional complexity, model fit was acceptable for all regions (CFI > 0.9, SRMR < 0.1). As an additional index of model validity and utility, we calculated the average variance explained (AVE) across indicators as 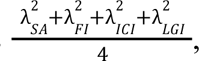 where *λ* refers to the factor loading of a given indicator.

Given the high degree of genetic overlap between regions, we calculated the number of effective tests for any subsequent regional analyses to more accurately correct for multiple comparisons among correlated tests. Here, we based our analysis on SA because it is the most widely studied indicator of cortical expansion and loads strongly onto the factor. We computed a 177×177 genetic correlation matrix encompassing the 177 regions of SA tested in this work. We estimated the nearest positive definite matrix using the Matrix package in R, and subsequently applied the Li and Ji method for estimating the number of effective tests^95^. For analyses conducted at the regional level, we used the estimated 51 independent tests for multiple comparisons corrections (*P* < 0.05/51 = 9.8e-4).

### Topographical annotation

To characterize the spatial organization of genetic influences on cortical expansion, we compared our empirical cortical maps to 10 canonical maps from the literature that index various aspects of brain organization. Specifically, these maps included:

1. the x-axis, reflecting the organization of cortical regions from lateral (outer) to medial (inner);
2. the y-axis, reflecting the organization of cortical regions from anterior (front) to posterior (back);
3. the z-axis, reflecting the organization of cortical regions from dorsal (upper) to ventral (lower);
4. the sensorimotor-association axis, indexing a gradient from unimodal, functionally specific primary sensorimotor regions to transmodal, functionally flexible association regions^20^;
5. a map of evolutionary expansion that indexes the degree of cortical expansion from macaques to humans^1,20^;
6. a map of developmental expansion that indexes the degree of cortical expansion during development^1^;
7. a gradient of anatomical hierarchy that reflects interregional patterns of T1w/T2w ratio, which approximates cortical myelination and laminar differentiation^20,96^;
8. the principal axis of functional connectivity, representing a unimodal-to-transmodal gradient^20,97^;
9. a gradient of plasticity that measures age-related changes in low-frequency fluctuation amplitude of resting-state fMRI data from eight to 22 years, where lower rank corresponds to reduced plasticity and higher rank corresponds to increased plasticity^98^; and
10. a map of excitation-inhibition balance, representing cross-sectional age-related differences in excitation-inhibition ratio based on structural and functional connectome data^21^.

For all canonical maps, regions were bilaterally averaged, matching our GWAS results^13^. For spins tests of enrichment and neuropsychiatric maps, we also included comparisons with an eleventh map: the local-global genetic similarity between regional and global cortical expansion (**Fig. 1e**).

To evaluate statistical significance, we used the neuromaps package to implement parcel-based spatial permutation spin tests with 10,000 permutations based on the Vázquez-Rodríguez method^23,99,100^. Spearman’s rank correlation was used, and two-sided p values were calculated. As our data were based on bilaterally averaged regions, we based our permutations on the left hemisphere. Here, we reported both nominal significance (*P_spin_* < 0.05) and corrected significance, as has been previously done in neuroimaging spatial permutation tests^20^. Since many of the canonical maps are correlated, we calculated the number of effective tests by estimating the nearest positive definite matrix using the Matrix package in R. We then used the Li and Ji method to estimate the number of effective tests^95^. We found that the 10 canonical maps corresponded to seven effective tests (*P* = 0.05/7 = 0.007), and eight effective tests (*P* = 0.05/8 = 0.006) when including an additional map of local-global similarity.

### Multivariate GWAS

We used Genomic SEM to conduct multivariate GWAS of the global cortical expansion factor. Here, we used unit loading identification when modeling SNP effects^101^, using SA to scale the factor. As recommended, data were filtered to those with imputation INFO scores > 0.6 and MAF > 0.01^16^ using the 1000 Genomes Phase 3 European as a reference panel^87^. Following these steps, 5,395,521 SNPs were available for analysis across SA, FI, ICI, LGI, and ICV. For each of these SNPs, we estimated their effect on the cortical expansion factor. We also calculated *Q*_SNP_, a heterogeneity statistic, to assess whether the estimated SNP effect plausibly operated through the common factor. For all downstream analyses (e.g., MiXeR, FUMA, MAGMA), we calculated the estimated sample size^101^ of the multivariate GWAS as 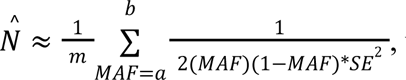 where *m* is the number of SNPs, and *a* and *b* reflect SNP MAF filters of 0.1 and 0.4, respectively.

For the relevant univariate and multivariate GWAS results, we used FUMA v1.8^25^ to identify the number of genomic risk loci and the number of lead SNPs. We used the default parameters, including genome-wide significance for lead variants (*P* < 5e-8), an *r^2^* threshold of 0.6 to identify independent significant SNPs, a second *r^2^* threshold of 0.1 to define lead SNPs, and LD block merging over 250kb. The 1000 Genomes Phase 3 European reference panel^87^ was used as an LD reference.

### Partitioned heritability and genetic covariance

We used Stratified Genomic SEM^35^ to test whether the genetic architecture of cortical expansion was enriched in functionally defined partitions of the genome. In brief, Stratified Genomic SEM extends Stratified LD Score regression^37^ to the multivariate context, testing whether shared and unique genetic signal across a set of phenotypes is enriched within particular genomic annotations. In the present study, we included 46 baseline annotations^36^, 45 of which were from the original Stratified LD Score regression authors^37^. We added a custom 46^th^ baseline annotation that indexed variation in loss-of-function intolerant genes^102^, which are defined as the 3,063 genes with probability of intolerance to heterozygous predicted-loss-of-function variation (pLI) greater than or equal to 0.9^41^. We also included 11 annotations indexing neurodevelopmental specificity as measured in BrainSpan^38^ and 12 annotations indexing cellular specificity in the human neocortex during the first trimester^39^. The neocortical cell-type expression data was previously processed via a public repository^40^ and downloaded via FUMA^25,40^. All annotations were created using the original LDSC Python package with a symmetrical 100kb window and the 1000 Genomes Phase 3 European reference panel^87^ as an LD reference. Accordingly, we corrected for 69 total annotations to assess significance for the global cortical expansion factor (*P* < 0.05/69 = 7.25e-4). For region-level analyses, which were only performed for select annotations, we corrected for the number of effective tests across brain regions (*P* < 0.05/51 = 9.8e-4).

For the BrainSpan and neocortical cell-type annotations, we sought to identify which genes were most specifically expressed in each developmental epoch or cell type. Using log-transformed reads-per-kilobase-million (BrainSpan) or counts-per-million (neocortical cell types), we first calculated the average expression of each gene across all categories within each dataset. Then, for each gene *i* and category *j*, we fit the regression model 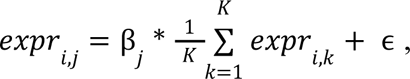 where *K* denotes the number of categories in the dataset: 11 developmental epochs for BrainSpan or 12 cell types for neocortical annotations. The resulting residuals therefore reflect a gene’s expression in a given category above or below the level expected based on that gene’s average expression across categories. We then formed annotations for each category based on the residualized expression values. Neurodevelopmental annotations were based on the top 10% of genes in each BrainSpan epoch, and the neocortical cellular annotations were based on the top 5% of genes in each cell type.

We conducted several additional sensitivity analyses. First, for the neocortical cell-type annotations, we repeated the Stratified Genomic SEM analyses while redefining annotations based on the top 2.5%, 5%, and 10% of residualized gene expression (**Figure S10**). Second, we evaluated the impact of defining annotations based on residualized expression values versus raw expression values (**Figure S11**).

### Gene-based association tests

To conduct a broader characterization of biological pathways implicated in cortical expansion, we performed gene-set enrichment analysis using MAGMA^29^ with the C5 subcollection from the Molecular Signatures Database^103,104^. Using MAGMA, we first conducted gene-based association analyses of 17,276 genes, applying a Bonferroni-corrected significance threshold (*P* = 0.05/17,276 = 2.89e-6). We then used these results in a gene-set enrichment analysis of 10,466 C5 gene sets. A total of 14 gene sets were excluded due to insufficient overlap (< 2 genes) with the GWAS results. A Bonferroni-corrected significance threshold was used to evaluate the significance of each tested gene set (*P* = 0.05/10,466 = 4.78e-6).

### Effector gene prioritization

We predicted the likely effector genes mediating locus-trait associations using FLAMES^32^. Briefly, FLAMES combines SNP-to-gene evidence and convergence-based evidence, outputting an estimated cumulative precision for each gene in fine-mapped loci. The first step of FLAMES involves fine-mapping. We identified approximately independent regions with significant signals (*P* < 5e-8) via FUMA with default parameters, as specified above. For each locus, we extended the boundary by ±100 kb and ran the SUm of SIngle Effects (SuSiE) model^28,105,106^ (susieR v0.14.2) to calculate posterior inclusion probability (PIP) for each variant and to identify 95% “credible sets” of putative causal variants. Default parameters were used, including a purity filter of 0.5 for credible sets. Subsequently, we used the FUMA v1.8^25^ implementation of MAGMA^29^ to calculate gene-based *Z* statistics. We applied the Polygenic Priority Score (PoPS)^107^ approach to these MAGMA results with the full annotation data to provide convergence-based evidence^108^.

The second step of FLAMES involves creating SNP-to-gene scores with an XGBoost classifier applied to the fine-mapped credible sets. Then, a final FLAMES score is calculated by linearly combining the raw XGBoost prediction scores and the locus-scaled PoPS scores. Cumulative precision is estimated based on a polynomial calibration curve that transforms scaled FLAMES scores (where all scaled scores in a locus sum to 1)^32^. As suggested by the developers, we considered effector genes as those with cumulative precision ≥ 0.75, and high-confidence effector genes as those with cumulative precision ≥ 0.999.

We tested whether the effector genes were enriched in genes previously linked to developmental disorders in the Deciphering Developmental Disorders (DDD) Study^33,34^. Here, we classified clinically relevant genes as those that had been implicated in developmental disorders with high confidence (**Table S9**). We used a background gene set that consisted of genes expressed in any GTEx v7 brain tissue downloaded from SynGO^109,110^. We then tested whether effector genes for cortical expansion were overrepresented in DDD genes with a one-sided Fisher’s exact test^109^.

### Genetic overlap with neuropsychiatric phenotypes

We evaluated patterns of genetic overlap between the cortical expansion factor and 20 neuropsychiatric phenotypes (**Table S13**) using Genomic SEM^35,111^ and MiXeR^51^. As noted previously, ICV was excluded from any analysis involving ALCP, ASB, CUD, OUD, or TUD due to data access stipulations; a comparable four-indicator cortical expansion factor model was used in these cases.

For Genomic SEM analyses, we used the *Q_Trait_* framework to test whether the associations between each external trait and the individual cortical expansion indicators were adequately explained by the common factor^111^. This involves the evaluation of two key metrics. First, we calculated the *Q_Trait_* statistic, which indexes the *χ*^2^ difference between the common pathway model, which estimates the association between the external trait and the common factor, and the independent pathways model, which estimates the association between the external trait and each indicator. Second, we evaluated the local standardized root mean squared residual (lSRMR) as an effect-size measure of heterogeneity, reflecting the degree to which indicator-level genetic correlations deviate from those implied by the common pathway model. For lSRMR to be considered indicative of heterogeneity, we required it to exceed two cutoffs: an absolute cutoff of lSRMR > 0.1 and a relative cutoff of lSRMR > 25% of the average root mean square genetic correlation. Significant and meaningful heterogeneity was only considered present if three criteria were met: (i) *Q*_Trait_ was significant after Bonferroni correction across traits, (ii) lSRMR was greater than the absolute cutoff, and (iii) lSRMR was greater than the relative cutoff. Based on these criteria, no neuropsychiatric traits exhibited heterogeneous associations with cortical expansion.

We also conducted bivariate MiXeR analyses to characterize potential patterns of concordant and discordant genomic overlap between cortical expansion and the neuropsychiatric phenotypes. We followed the same procedures as outlined in the analysis of FI, ICI, ICV, LGI, and SA. Six neuropsychiatric traits (ASB, ANX, PAN, OCD, TS, SRD) did not exhibit adequate univariate model fit and were excluded from bivariate MiXeR analysis (**Table S11**). All bivariate analyses between cortical expansion and neuropsychiatric phenotypes exhibited stable model fits across random iterations (**Fig. S20**).

## Acknowledgements

This study was supported by the National Human Genome Research Institute (M.R.: 5T32HG010464) and the National Institute of Mental Health (T.T.M.: K08MH135343). The present analyses would not have been possible without the enormous efforts put forth by the investigators and participants from the ENIGMA Consortium, iPSYCH, Million Veteran Program, PsychENCODE, Psychiatric Genomics Consortium, UK Biobank, and related cohorts.

## Author Contributions

T.T.M. and M.R. conceptualized the study. T.T.M. and J.W.S. supervised the study. M.R. led the analysis with assistance from T.T.M., M.V., and C.M.W. T.T.M. and M.R. prepared the figures. T.T.M., M.R., and H.F.J. prepared supplementary information. All authors contributed to and critically reviewed the manuscript.

## Competing Interests

J.W.S. is a member of the Scientific Advisory Board of Sensorium Therapeutics and holds stock options; has received consulting fees from Tempus, Inc.; and has received grant support from Biogen, Inc. All other authors declare no competing interests.

## Data Availability

GWAS data for SA, FI, ICI, and LGI were downloaded from: https://portal.camide.cam.ac.uk/overview/483. GWAS data for ICV were downloaded from the ENIGMA website https://enigma.ini.usc.edu/research/download-enigma-gwas-results/. Download links for the neuropsychiatric GWAS data are detailed in Table S13. The LD scores and HapMap3 reference data used in the EUR-like Genomic SEM analyses were downloaded from: https://github.com/GenomicSEM/GenomicSEM/wiki. C5: Ontology gene sets were downloaded from the Molecular Signatures Database: https://www.gsea-msigdb.org/gsea/msigdb/collections.jsp The neocortical cell type RNA sequencing data were downloaded from FUMA: https://fuma.ctglab.nl/downloadPage and originally processed at: https://github.com/tanyaphung/scrnaseq_viewer/. Developmental epoch RNA sequencing data were downloaded from BrainSpan: https://www.brainspan.org/static/download. Gene-level loss-of-function data were downloaded from gnomAD: https://gnomad.broadinstitute.org/downloads. Baseline annotations used in Stratified Genomic SEM were downloaded from: https://github.com/GenomicSEM/GenomicSEM/wiki/6.-Stratified-Genomic-SEM. Genes implicated in developmental disorders were downloaded from the Gene2Phenotype site:

https://www.ebi.ac.uk/gene2phenotype/panel/DD. Background gene sets that consisted of genes expressed in any GTEx v7 brain tissue downloaded from SynGO: https://www.syngoportal.org/download.

## Code Availability

We used several publicly available pieces of software in our study. Genomic SEM v0.0.5 (https://github.com/GenomicSEM/) was used in LDSC, Genomic SEM, Stratified Genomic SEM, and multivariate GWAS analyses. MAGMA v1.10 (https://cncr.nl/research/magma/) was used for evaluating enrichment in the C5:GO gene sets. The LDSC package was used for annotation creation for Stratified Genomic SEM (https://github.com/bulik/ldsc) MiXeR v1.3 (https://github.com/precimed/mixer/tree/f56a44bde5bd3f8bc0d143d89d1b629035b2d675) was used to calculate polygenic overlap. Neuromaps 0.0.5 (https://netneurolab.github.io/neuromaps) was used for spin tests. FUMA v1.8 (http://fuma.ctglab.nl/) was used for identifying genomic loci. susieR v0.14.2 (https://stephenslab.github.io/susieR/) was used for fine-mapping as input into FLAMES. PoPs (https://github.com/FinucaneLab/pops) was used to calculate polygenic priority scores as input into FLAMES. FLAMES (https://github.com/Marijn-Schipper/FLAMES) was used to predict effector genes.

