## Supplementary Text and Figures for "Mapping the genetic architecture of human cortical expansion and its links to neuropsychiatric disorders"

### Supplementary Information

### *S1. Imaging MiXeR results*

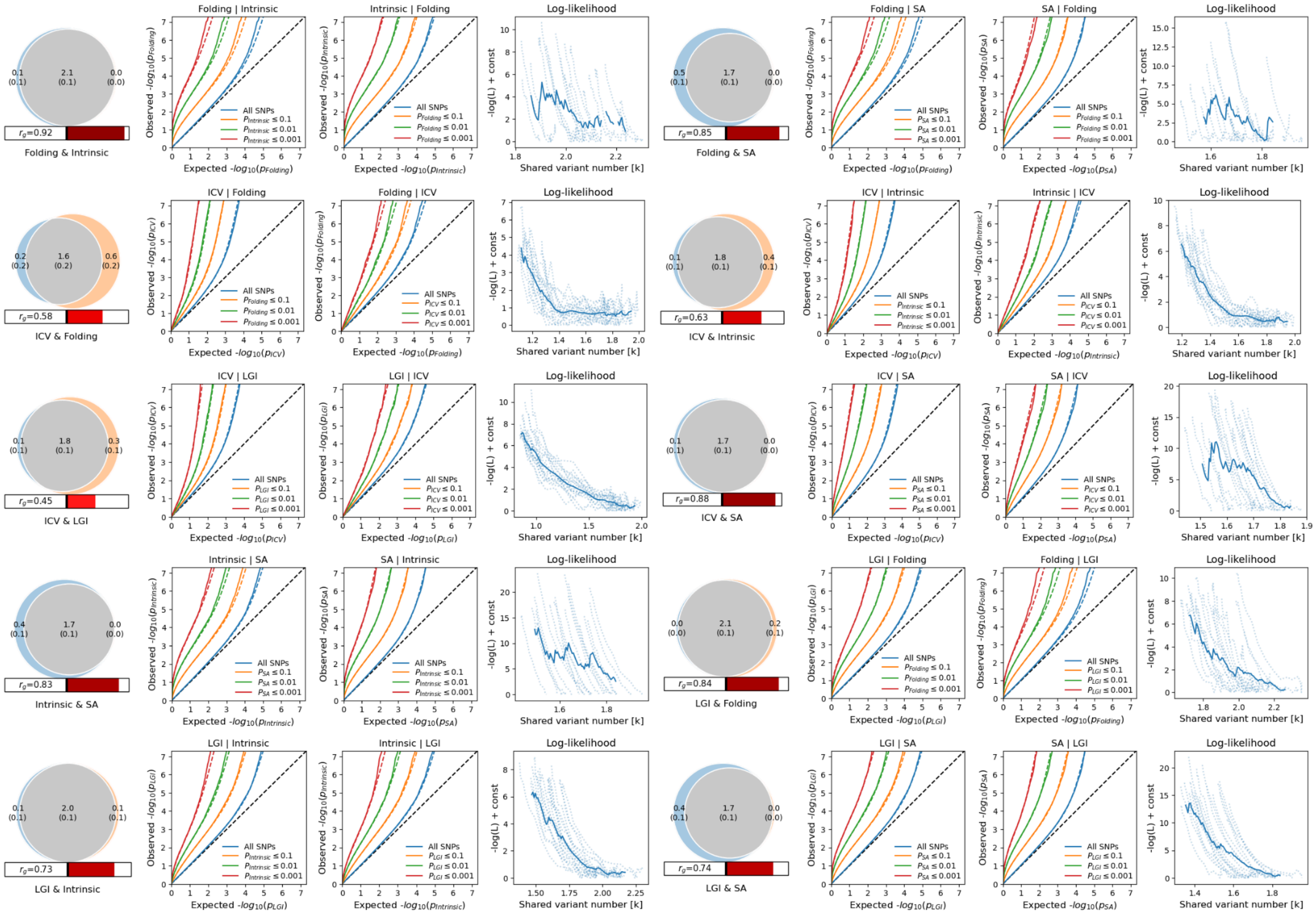


**Supplementary Figure S1**. **Bivariate MiXeR fits for cortical expansion indicators.** For each pair of five imaging traits, their genetic overlap (number of causal variants at 90% SNP heritability) is shown as a Venn diagram. In addition, conditional QQ plots show the signal of one GWAS stratified by GWAS signal for the other trait. Finally, log-likelihood plots demonstrate the fit of the model across 20 random iterations, each selecting a random subset of 600,000 SNPs.

***S2. Structural equation models of cortical expansion***

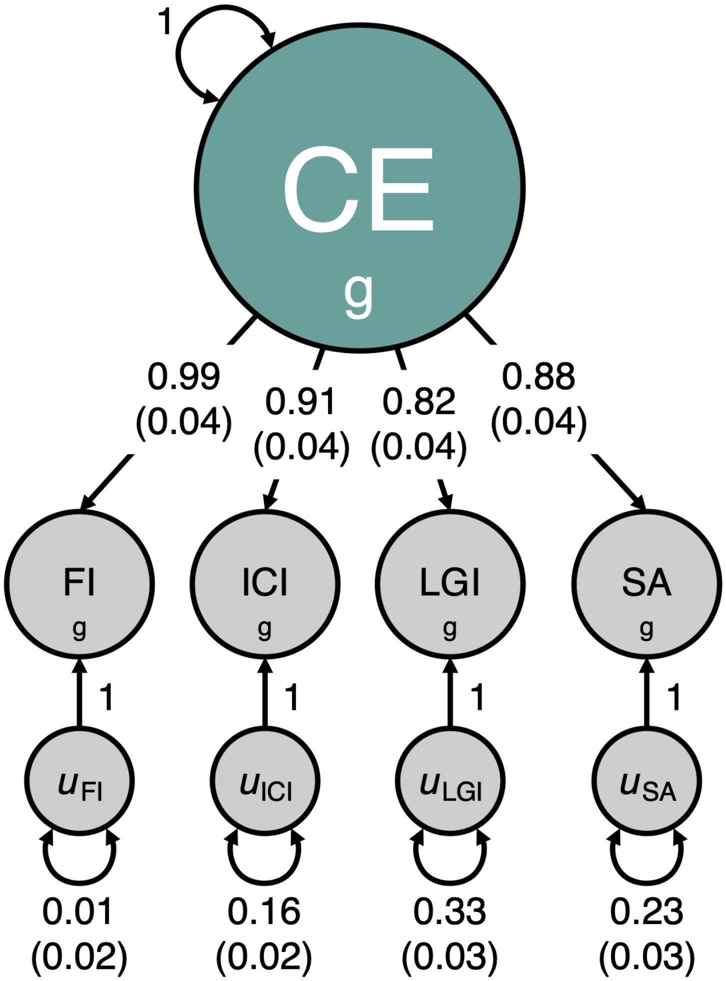


**Supplementary Figure S2. Common factor model path diagram without ICV**. Structural equation model path diagram for a global cortical expansion model. Standardized loadings are shown across the four indicators. FI: folding index; ICI: intrinsic curvature index; LGI: local gyrification index; SA: surface area.

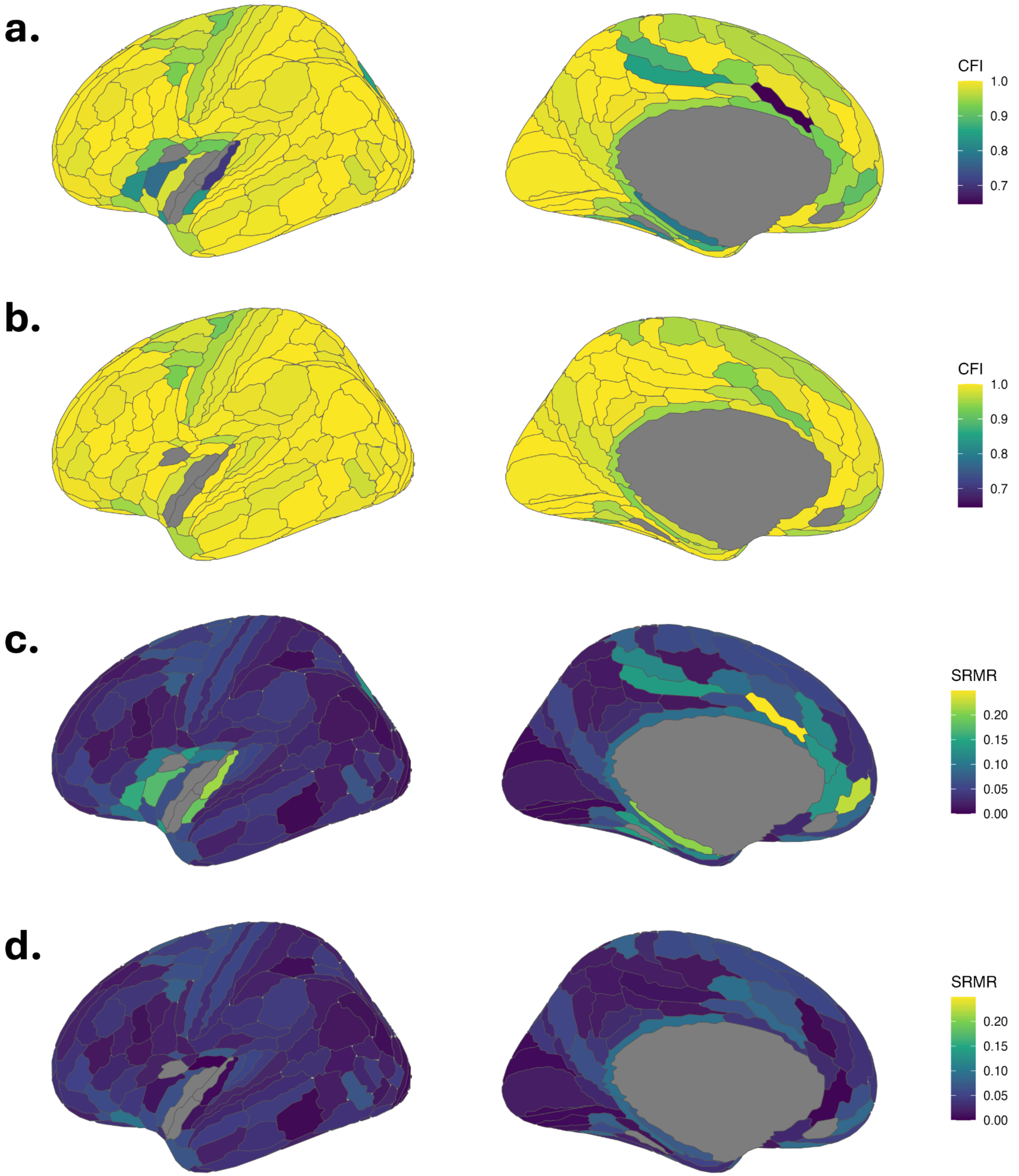


**Supplementary Figure S3**. **Model fit evaluated across brain regions.** Common factor analysis was performed for 177 bilateral regions with four regional indicators of cortical expansion: surface area, folding index, intrinsic curvature index, and local gyrification index. While the HCP atlas has 180 regions, three regions (52, PHA2, PI) were excluded due to negative or non-significant heritability for FI and ICI, leaving 177 regions. For an additional three regions (FOP3, s32, PoI1), heritability for FI was not significant, so this term was excluded from the model. Without FI, the model was saturated and thus interpreting model fit was not meaningful, leaving 174 regions to interpret model fit. The Comparative Fit Index (CFI) and Standardized Root Mean Squared Residual (SRMR) were calculated as measures of fit across these 174 bilateral regions, at baseline (**a, c**) and after accounting for additional residual covariance between the two indicators with the highest residual covariance (**b, d**; ICI and FI for 17 regions, and SA and LGI for five regions).

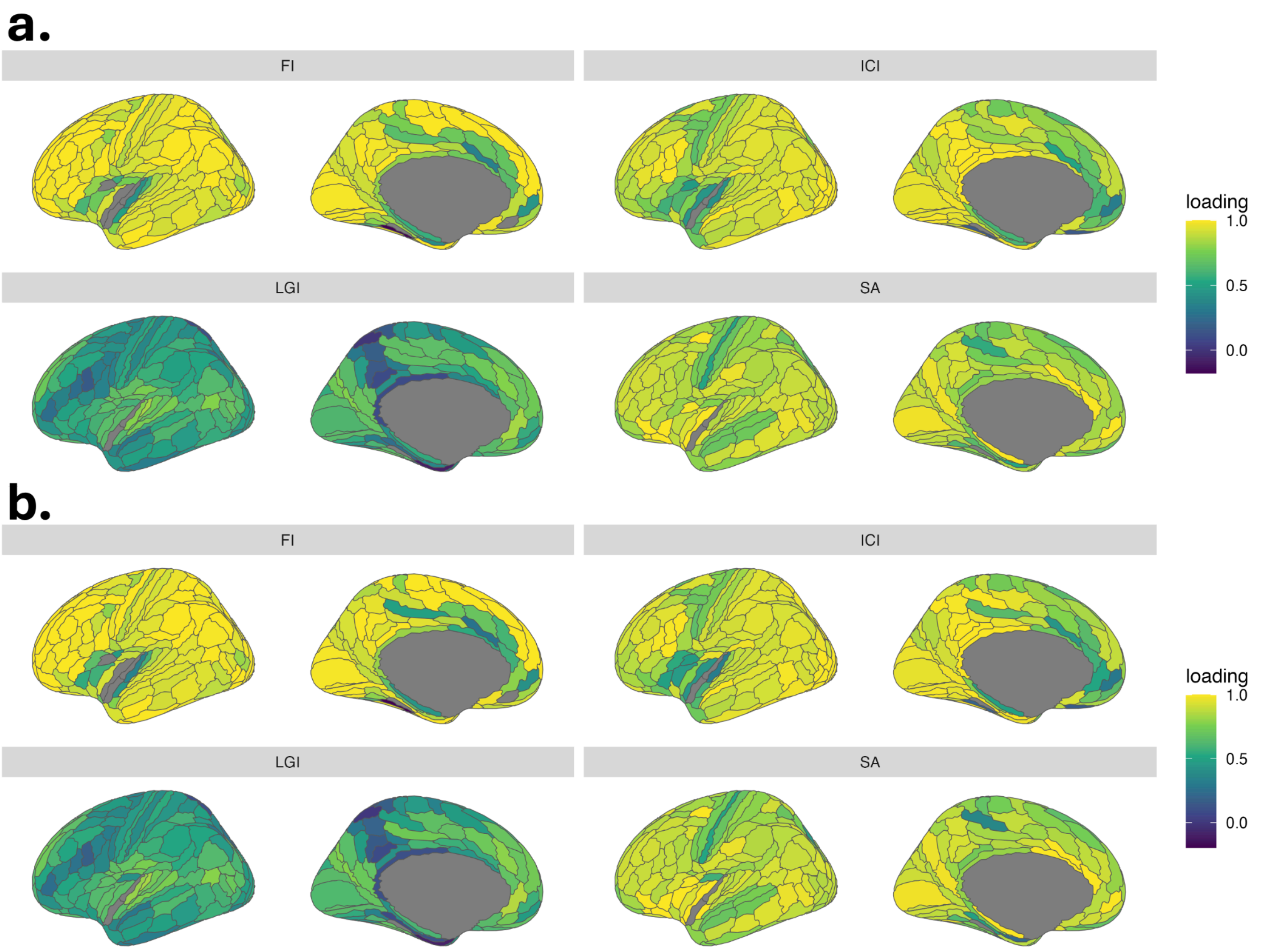


**Supplementary Figure S4**. **Regional loadings of model fits.** Standardized regional loadings are shown for **a)** the baseline model, and **b)** after correcting for residual covariance to achieve comparative fit index (CFI) > 0.9 and standardized root mean square residual (SRMR) < 0.1.

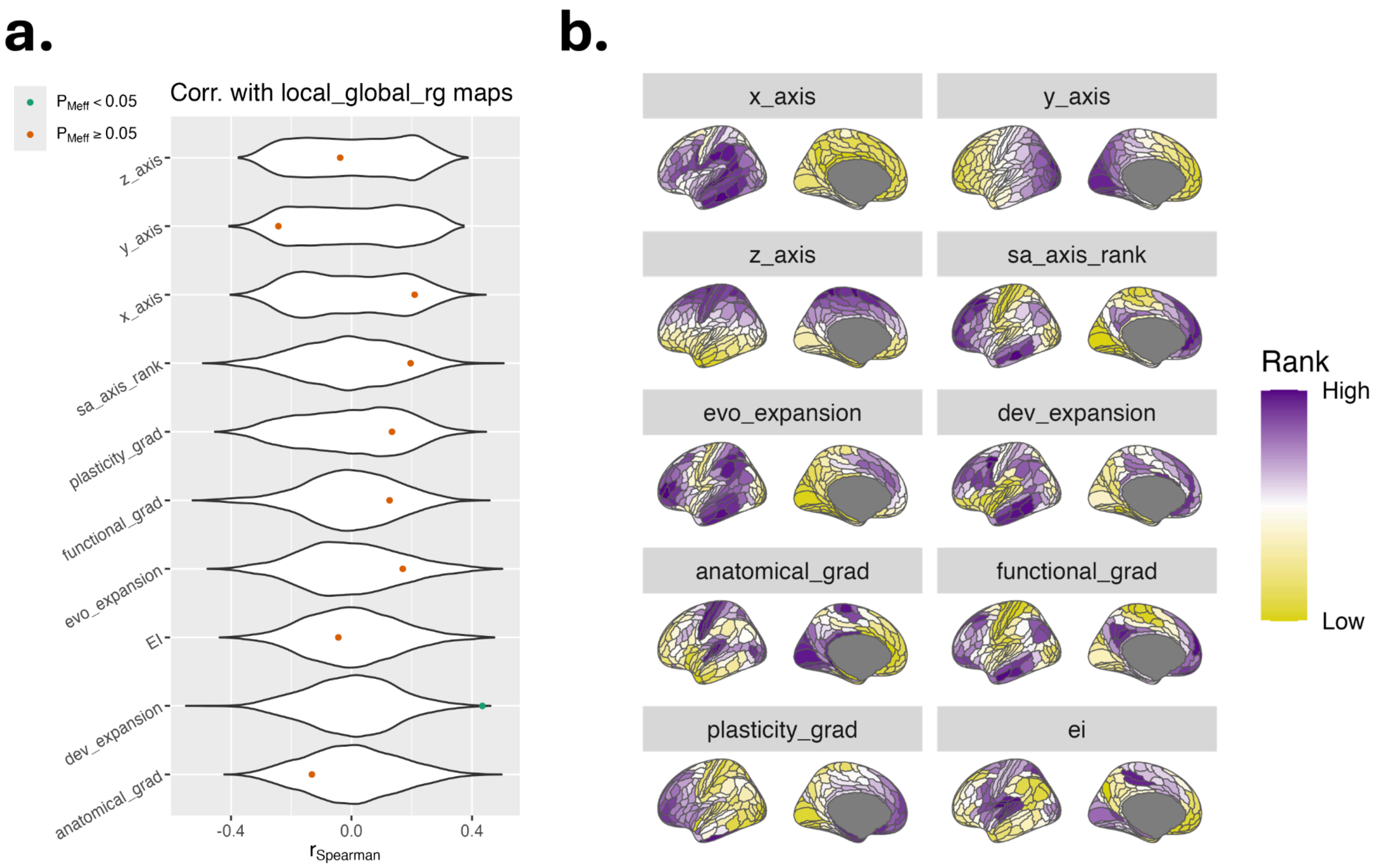


**Supplementary Figure S5**. **Spatial correlation between local-global correlation maps and existing anatomical/functional maps. a)** Results of a spin test with 10,000 permutations. Significance is shown after correcting for the number of effective tests (seven). **b)** Plots of canonical maps. **Map definitions:** X: Lateral-medial axis; reflects the organization of cortical regions from lateral (outer) to medial (inner). Y: Anterior-posterior axis; reflects the organization of cortical regions from anterior (frontal) to posterior (back). Z: dorsal-ventral axis; reflects the organization of cortical regions from dorsal (upper) to ventral (lower). S-A ranks: A multimodal sensorimotor-association gradient; indexes transitions from primary sensorimotor areas, which are responsible for basic sensory and motor functions, to higher-order association cortices involved in complex cognitive processes[1]. Evolutionary expansion: Evolutionary hierarchy; reflects the degree of cortical expansion observed in humans compared to non-human primates, highlighting regions that have expanded significantly during evolution [2]. Developmental expansion: Developmental hierarchy; represents the degree of regional cortical expansion observed in humans across development, where some regions expand significantly during childhood and adolescence and others less so[2]. Anatomical gradient: T1w/T2w, a measure of structural hierarchy that approximates cortical myelination and laminar differentiation[1,3]. Functional gradient: Functional hierarchy; a unimodal-to-transmodal gradient that reflects the organization of the cortex from regions specialized for specific functions to regions involved in multiple integrative functions[1,4]. Cortical plasticity gradient: measures age-related changes in low-frequency fluctuation amplitude of resting-state fMRI data from eight to 22 years, thought to represent changes in plasticity (low rank: decreasing plasticity, high rank: increasing plasticity)[5]. EI: Excitation-inhibition ratio; the excitation-inhibition map represents cross-sectional age-related differences in excitation-inhibition ratio, based on structural and functional connectome data[6].

#### *S3. $Q_{SNP}$ analyses*

The signal due to cortical expansion was stronger than that of $Q_{SNP}$ (**Fig. S6**). The cortical expansion signal exhibited substantial inflation ($\lambda_{GC}$=1.26, mean $\chi^2$=1.35), with limited effects of population stratification or other confounding (LD score regression intercept=1.02 [SE=0.01], attenuation ratio=0.06 [SE=0.03]). The $Q_{SNP}$ signal exhibited substantial inflation ($\lambda_{GC}$=1.30, mean $\chi^2$=1.32), with limited effects of population stratification or other confounding (LD score regression intercept=1.01 [SE=0.01], attenuation ratio=0.04 [SE=0.03]).

To test whether the FUMA or MAGMA results were sensitive to heterogeneity, we applied a model to reduce $Q_{SNP}$. For all SNPs with significant $Q_{SNP}$ (P<5e-8), we adjusted the model used to form multivariate summary statistics. As an initial step in the adjustment, we tested five separate models, adding a direct pathway between the SNP and each indicator (one per model). Then, we selected the indicator which reduced $Q_{SNP}$ the most and retained those results. FUMA identified 45 genomic loci after this correction, as opposed to 48 originally. We overlaid the original genomic loci on the updated Manhattan plot (**Fig. S7**). MAGMA identified the same 122 genes as significantly associated with CE. However, MAGMA gene set enrichment analysis identified six of the seven gene sets as significant (**Table S6**). The missing gene set was related to positive regulation of hydrogen peroxide mediated programmed cell death.

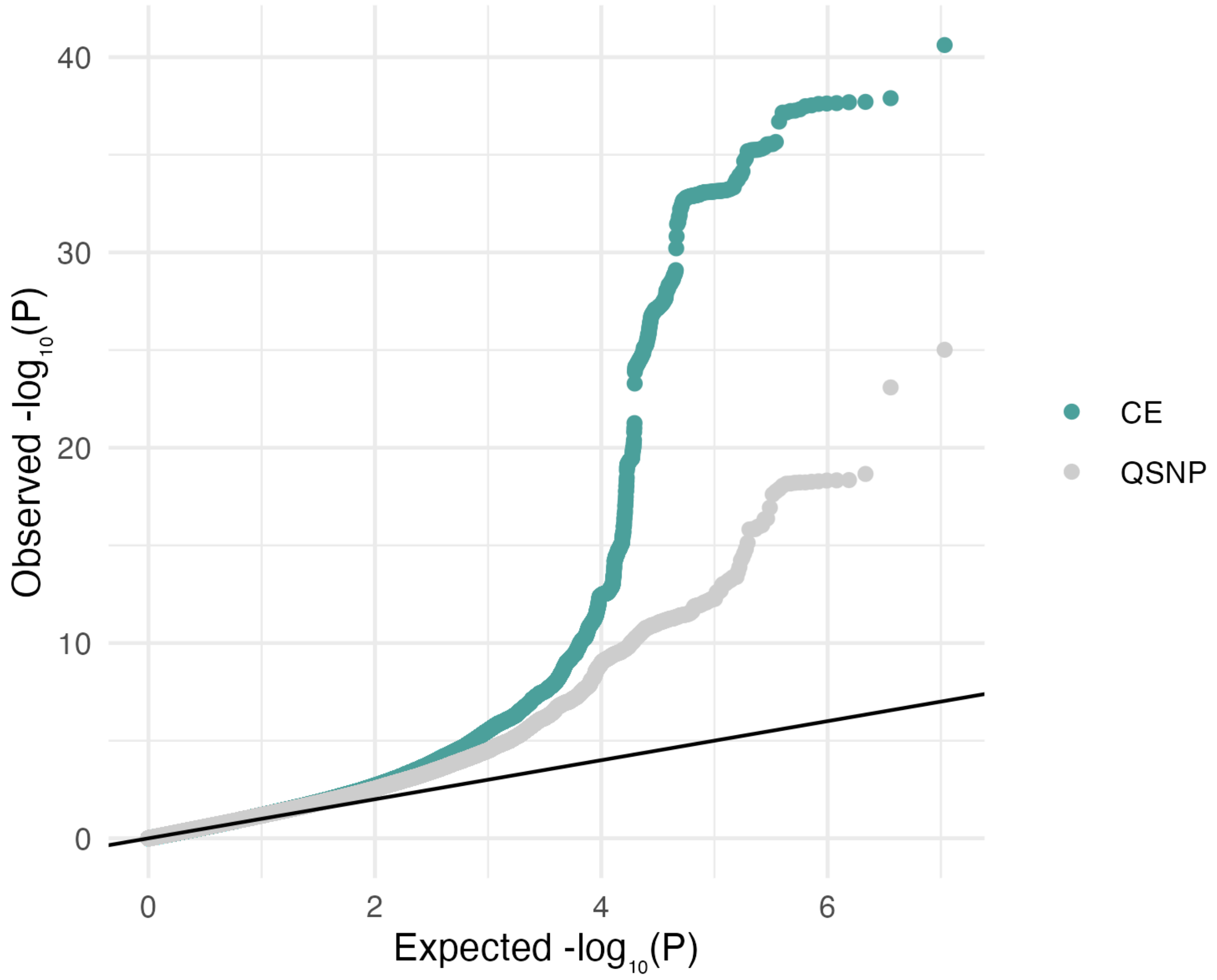


**Supplementary Figure S6. QQ-plot for cortical expansion factor GWAS and $Q_{SNP}$ signal.** For global cortical expansion, we compared the multivariate GWAS signal (cortical expansion: CE) with the heterogeneity signal ($Q_{SNP}$), demonstrating increased CE signal relative to its heterogeneity across indicators.

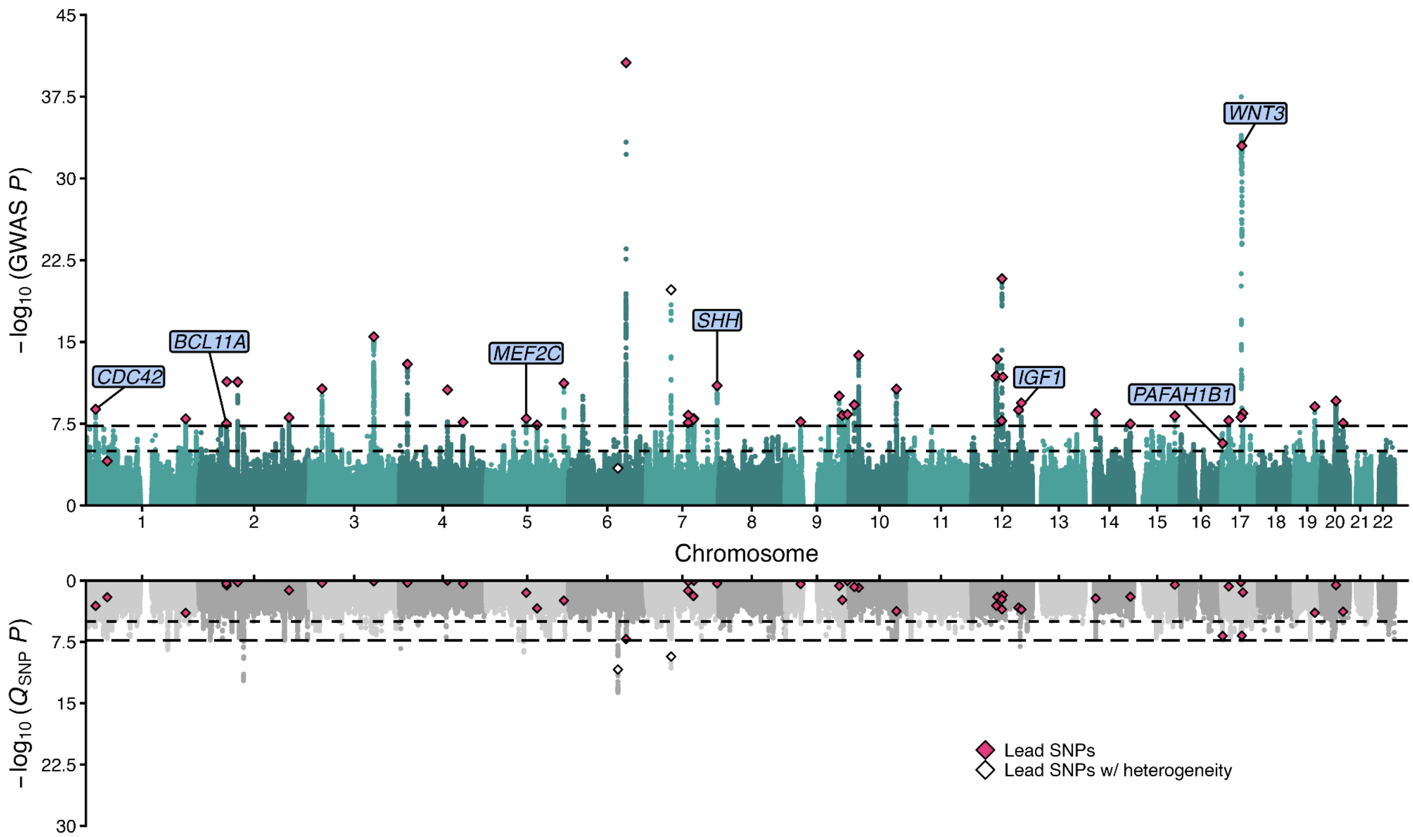


**Supplementary Figure S7. QQ-plot for cortical expansion factor GWAS and $Q_{SNP}$ signal after $Q_{SNP}$ correction.** $Q_{SNP}$ correction was applied to SNPs with significant $Q_{SNP}$ by adding a direct pathway between the SNP and the indicator which reduced $Q_{SNP}$ the most. Diamonds represent lead SNPs for loci identified by FUMA for the uncorrected GWAS (**Fig. 2**). Annotated genes include seven effector genes (cumulative precision $\geq 0.75$) that were identified via FLAMES in the uncorrected GWAS (**Fig. 2**) and overlap with the Deciphering Developmental Disorders Study.

#### *S4. Stratified Genomic SEM sensitivity analyses*

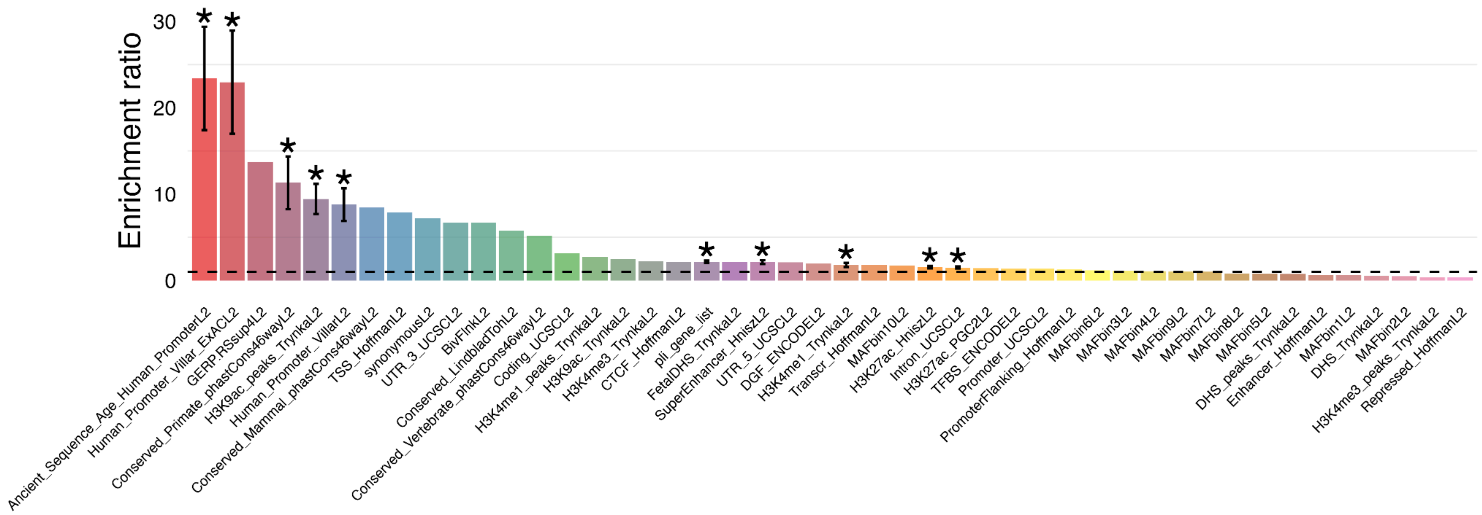


**Supplementary Figure S8. Enrichment in baseline annotations.** Stratified Genomic SEM was used to assess enrichment in 45 baseline annotations[7] from the original S-LDSC authors[8], as well as a 46th annotation, reflecting loss-of-function intolerant genes, which are defined as the 3,063 genes with probability of intolerance to heterozygous predicted-loss-of-function variation (pLI) greater than or equal to 0.9[9]. The error bars reflect the standard error of the enrichment estimate. Significance was assessed with a Bonferroni correction across 69 annotations tested in Stratified Genomic SEM ($P<0.05/46=7.25e-4$).

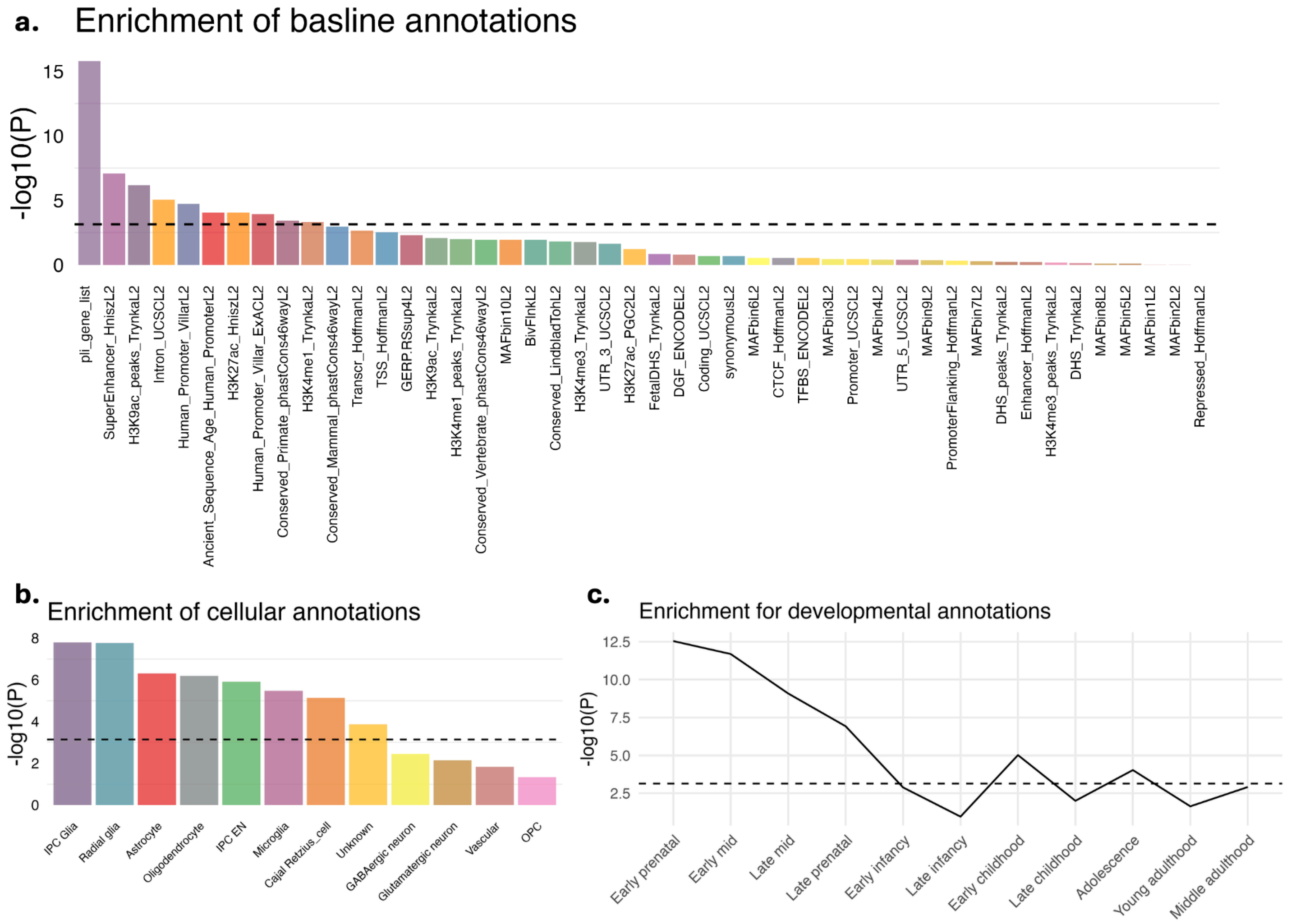


**Supplementary Figure S9**. **P values for Stratified Genomic SEM enrichment analyses.** P values for enrichment were analyzed for **a)** baseline annotations, **b)** neocortical cell type annotations, and **c)** developmental annotations. Significance was assessed with a Bonferroni correction across all 69 annotations, and the significance threshold is marked by a dashed horizontal line.

We performed a sensitivity analysis to evaluate enrichment of neocortical cell type annotations when considering the top 2.5%, 5%, and 10% of genes to form an annotation (**Fig. S10**). Unlike BrainSpan, for which we used the top 10% of genes in an annotation to capture broad developmental patterns, we sought to capture more specific cellular signals for the cell type annotations. Using an annotation size of 5% gave us the greatest specificity of enrichment.

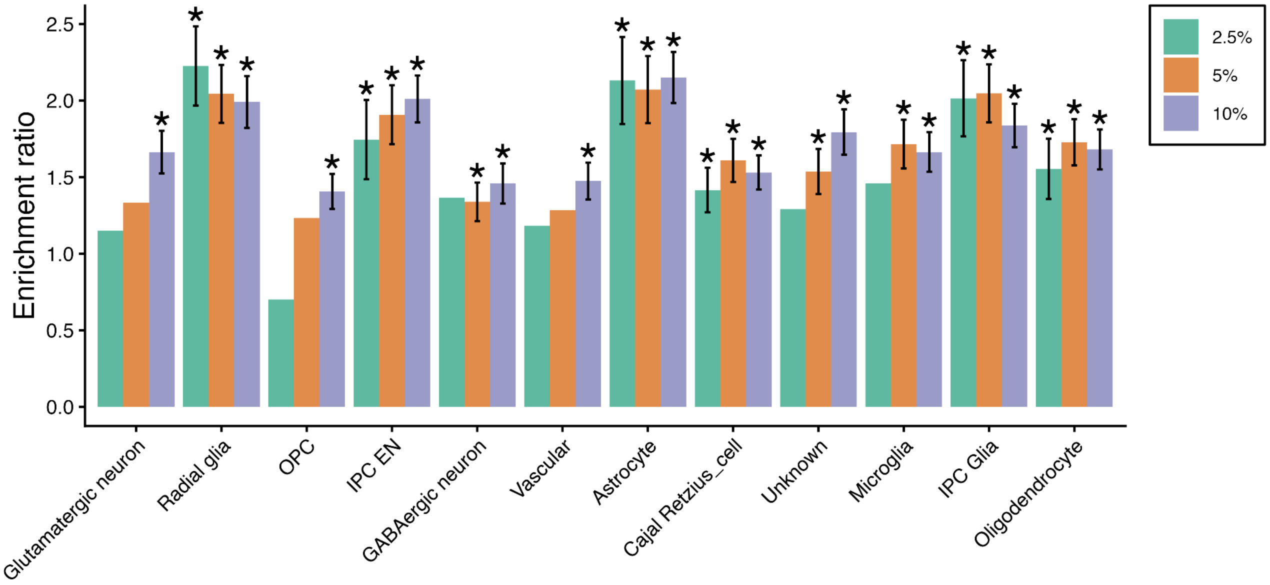


**Supplementary Figure S10. Neocortical cellular annotation size sensitivity analysis**. Comparison of the enrichment of cortical expansion in annotations when taking the top 2.5%, 5%, and 10% of residualized gene expression. Error bars reflect the standard error of the enrichment estimate.

An additional sensitivity analysis forming annotations based on the top 10% raw gene expression demonstrated similar trends but reduced specificity of enrichment relative to the residualized approach (**Fig. S11**).

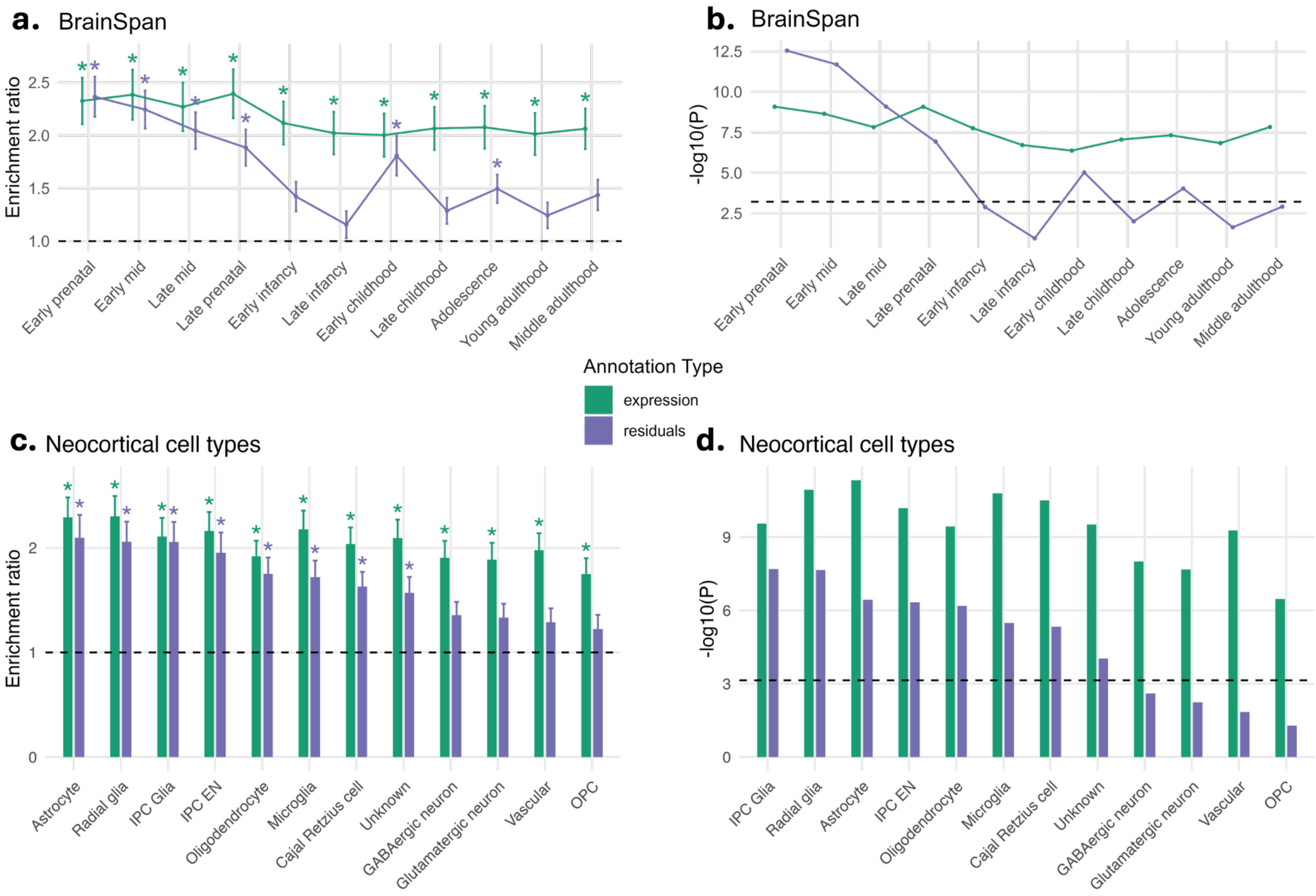


**Supplementary Figure S11**. **Comparison of residualized and raw expression annotation approaches for Stratified Genomic SEM.** The **a)** enrichment ratio and **b)** P value of the enrichment ratio are shown for BrainSpan, as well as for neocortical cell type annotations (**c, d**). In the raw approach, the top 10% of genes with the highest raw expression were selected to form an annotation. In the residualized approach, the top 10% of genes with the highest expression after regressing out the average expression across all annotations were selected. Error bars reflect the standard error of the enrichment estimate.

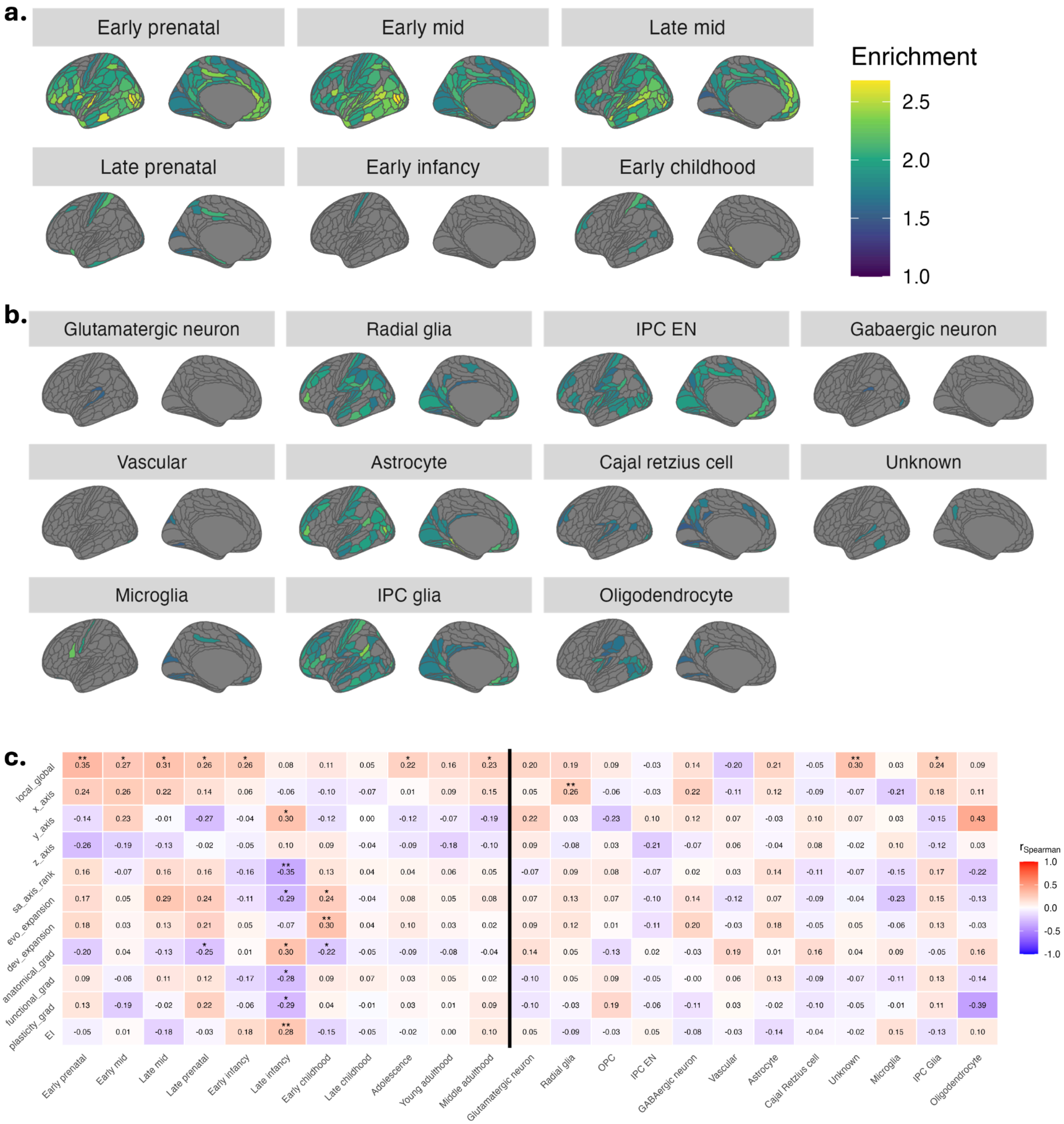


**Supplementary Figure S12. Characterizing enrichment of the cortical expansion factor. a)** Regional enrichment for developmental annotations. **b)** Regional enrichment for neocortical cell type annotations. For both **a)** and **b)**, the significance threshold is defined by 51 effective tests (P=0.05/51=9.80e-4), derived from the genetic correlation structure of the 177 regions. **c)** Correlation between anatomical-functional maps and developmental (left) and neocortical cell type (right) enrichment maps. A single asterisk (*) reflects $P_{spin}<0.05$, and a double asterisk (**) reflects significance corrected for the effective number of tests (see descriptions in Methods). **Map definitions**[1]**:** Y: Anterior-posterior axis; reflects the organization of cortical regions from anterior (frontal) to posterior (back). Z: dorsal-ventral axis; reflects the organization of cortical regions from dorsal (upper) to ventral (lower). S-A ranks: A multimodal sensorimotor-association gradient; indexes transitions from primary sensorimotor areas, which are responsible for basic sensory and motor functions, to higher-order association cortices involved in complex cognitive processes[1]. Evolutionary expansion: Evolutionary hierarchy; reflects the degree of cortical expansion observed in humans compared to non-human primates, highlighting regions that have expanded significantly during evolution [2]. Developmental expansion: Developmental hierarchy; represents the degree of regional cortical expansion observed in humans across development, where some regions expand significantly during childhood and adolescence and others less so[2]. Anatomical gradient: T1w/T2w, a measure of structural hierarchy that approximates cortical myelination and laminar differentiation[1,3]. Functional gradient: Functional hierarchy; a unimodal-to-transmodal gradient that reflects the organization of the cortex from regions specialized for specific functions to regions involved in multiple integrative functions [1,4]. Cortical plasticity gradient: measures age-related changes in low-frequency fluctuation amplitude of resting-state fMRI data from eight to 22 years, thought to represent changes in plasticity (low rank: decreasing plasticity, high rank: increasing plasticity)[5]. EI: Excitation-inhibition ratio; the excitation-inhibition map represents cross-sectional age-related differences in excitation-inhibition ratio, based on structural and functional connectome data[6].

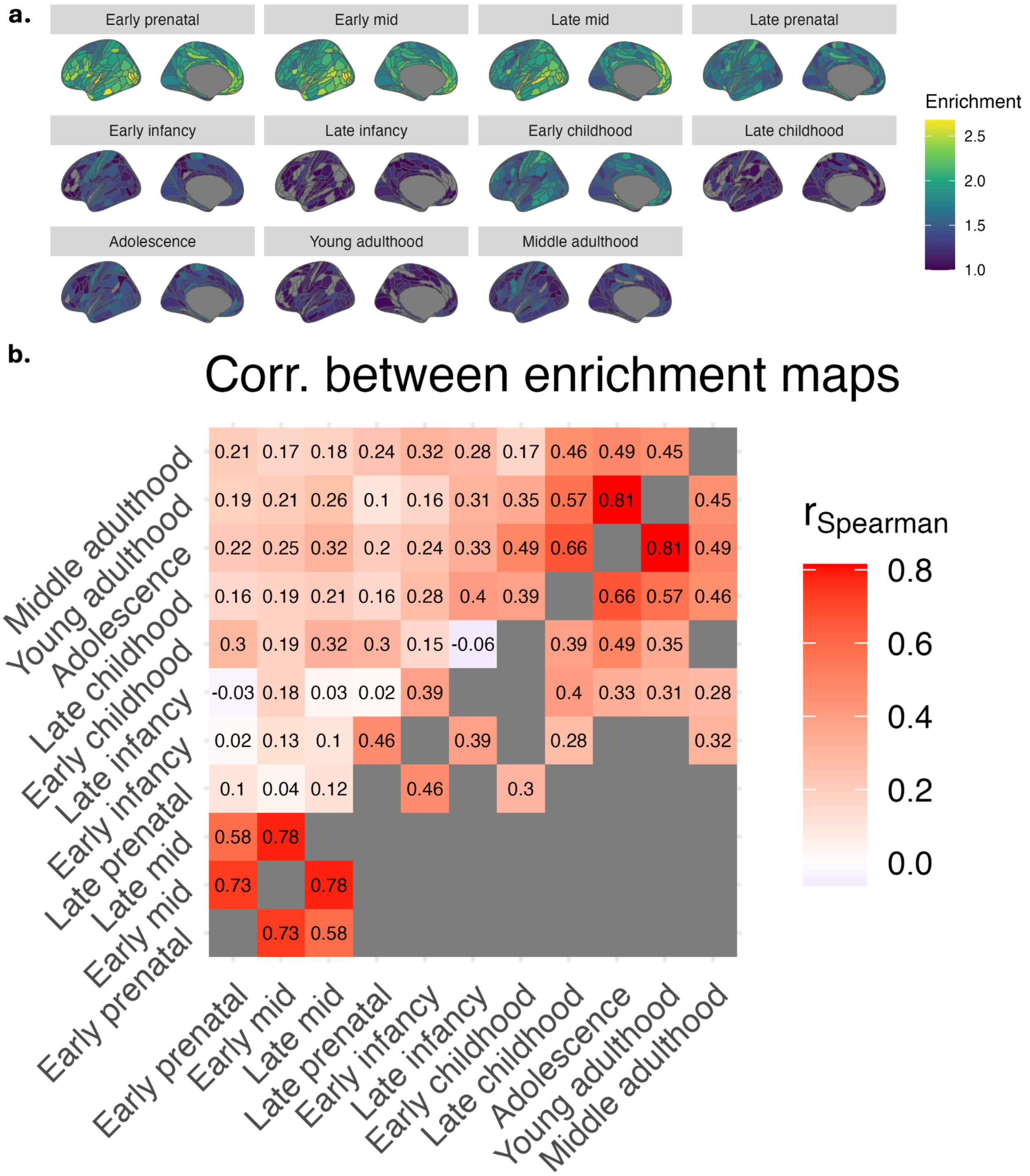


**Supplementary Figure S13. Comparison of developmental enrichment maps of cortical expansion. a)** The unthresholded enrichment ratios maps are presented for each of the 11 developmental epochs. Gray regions reflect poor fit in stratified analyses. **b)** Spatial correlation between unthresholded enrichment maps, where the upper triangle shows all correlations, and the lower triangle shows significant correlations derived from spin tests ($P = 0.05 / 55$ pairwise tests = 9.09e-4).

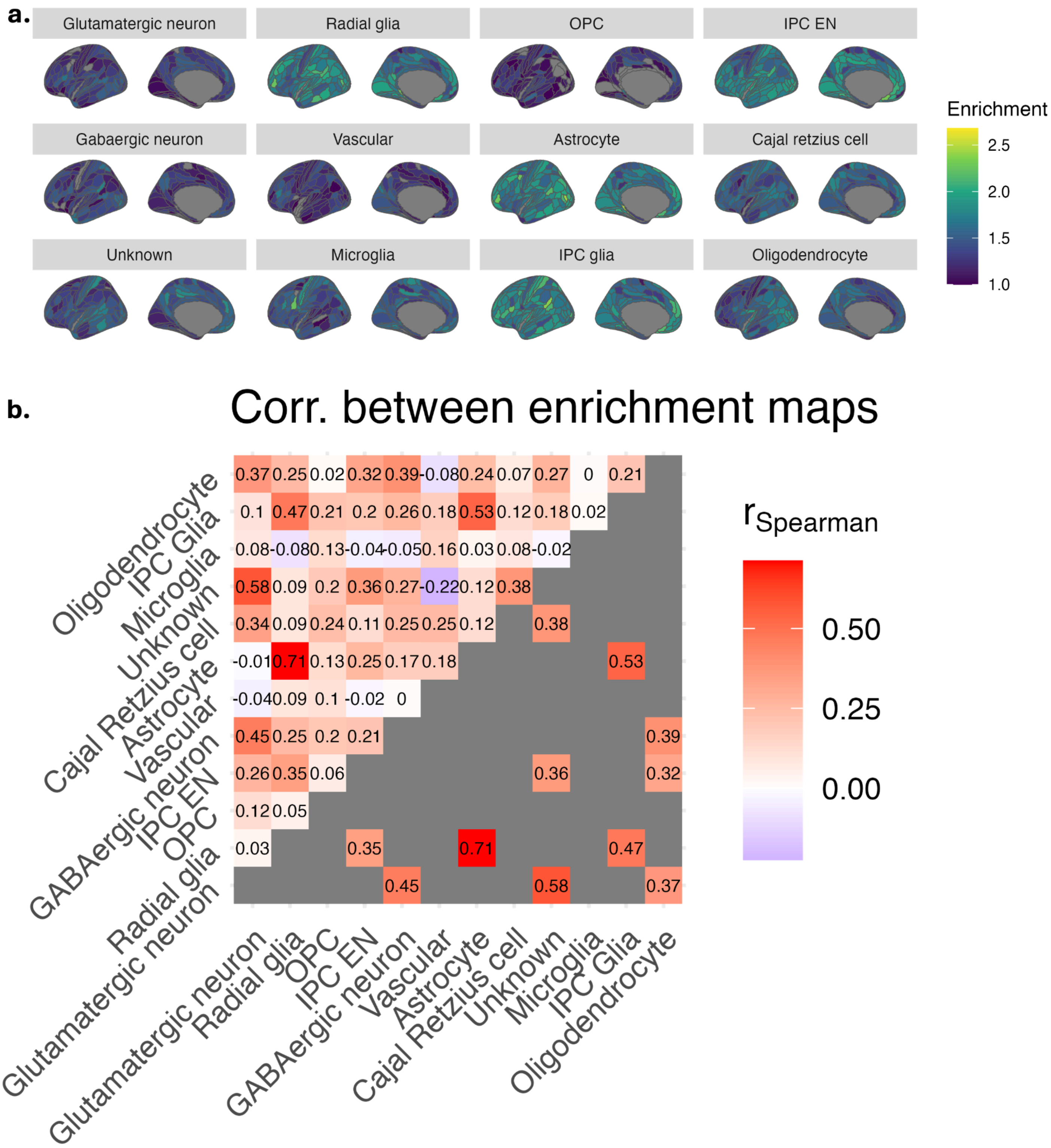


**Supplementary Figure S14. Comparison of neocortical cell type enrichment maps of cortical expansion. a)** The unthresholded enrichment ratios maps are presented for each of the 12 neocortical cell types. **b)** Spatial correlation between unthresholded enrichment maps, where the upper triangle shows all correlations, and the lower triangle shows significant correlations derived from spin tests ($P$ = 0.05 / 66 pairwise tests = 7.58e-4).

***S5. External correlations sensitivity analyses***

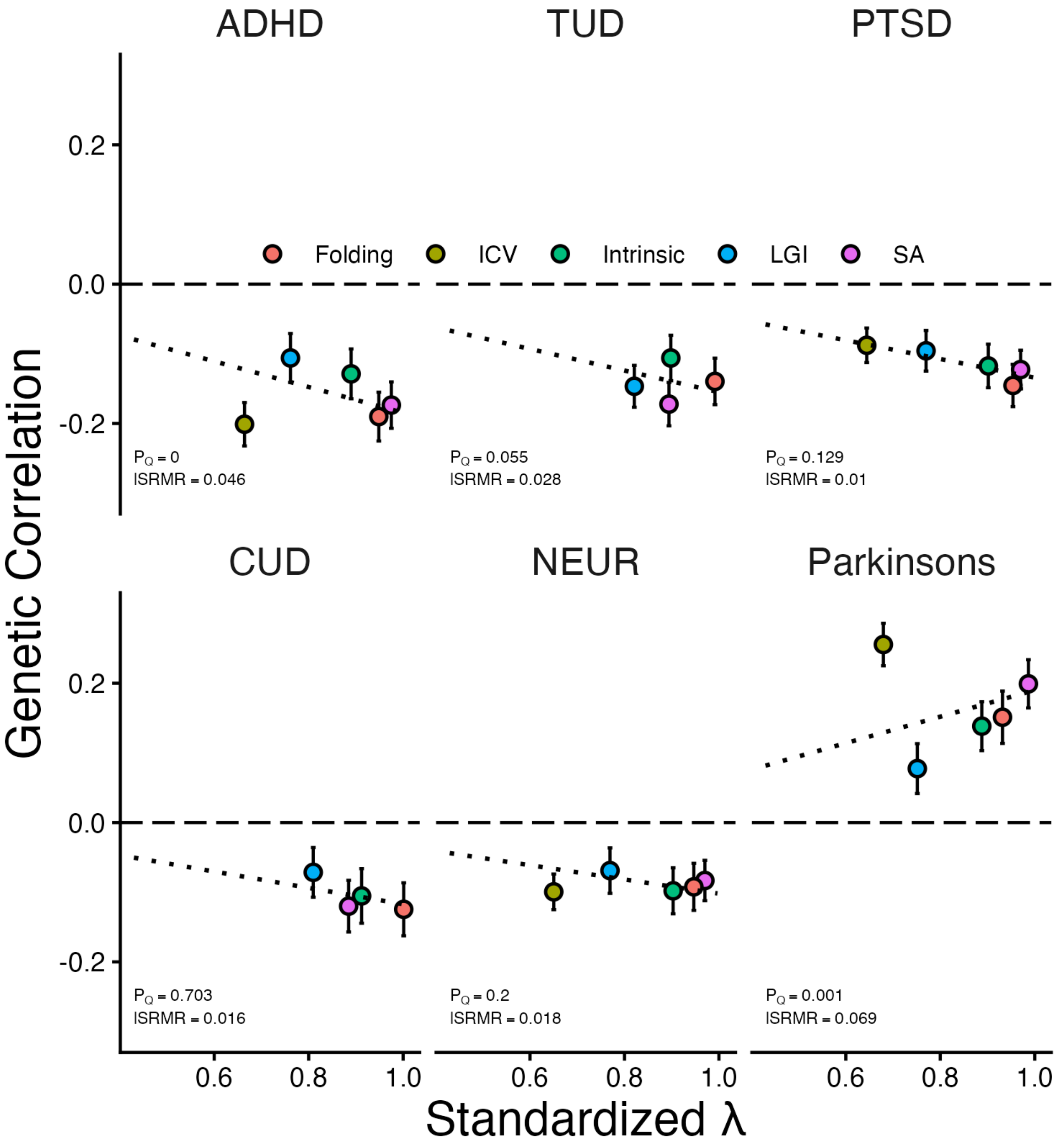


**Supplementary Figure S15. $Q_{Trait}$ analyses for external correlations.** $Q_{trait}$ analysis to determine whether the cortical expansion factor explains the relationship between individual imaging indicators and the psychiatric traits/disorders. The horizontal axis shows the loadings of each indicator, while the vertical axis shows the correlation of each indicator with the external trait. Deviations from the line of best fit show indicators that drive higher $Q_{Trait}$. Significant and meaningful heterogeneity was only considered present if three criteria were met: 1) $Q_{Trait}$ is significant after Bonferroni correction across traits 2) local standardized root mean squared residual (lSRMR) is greater than the absolute cutoff of 0.1, and 3) lSRMR is greater than the relative cutoff of 25% of the average root mean square genetic correlation[10]. The error bars represent the standard error of the genetic correlation estimate. For traits that are considered sensitive (CUD, TUD), ICV was excluded from the cortical expansion structural equation model.

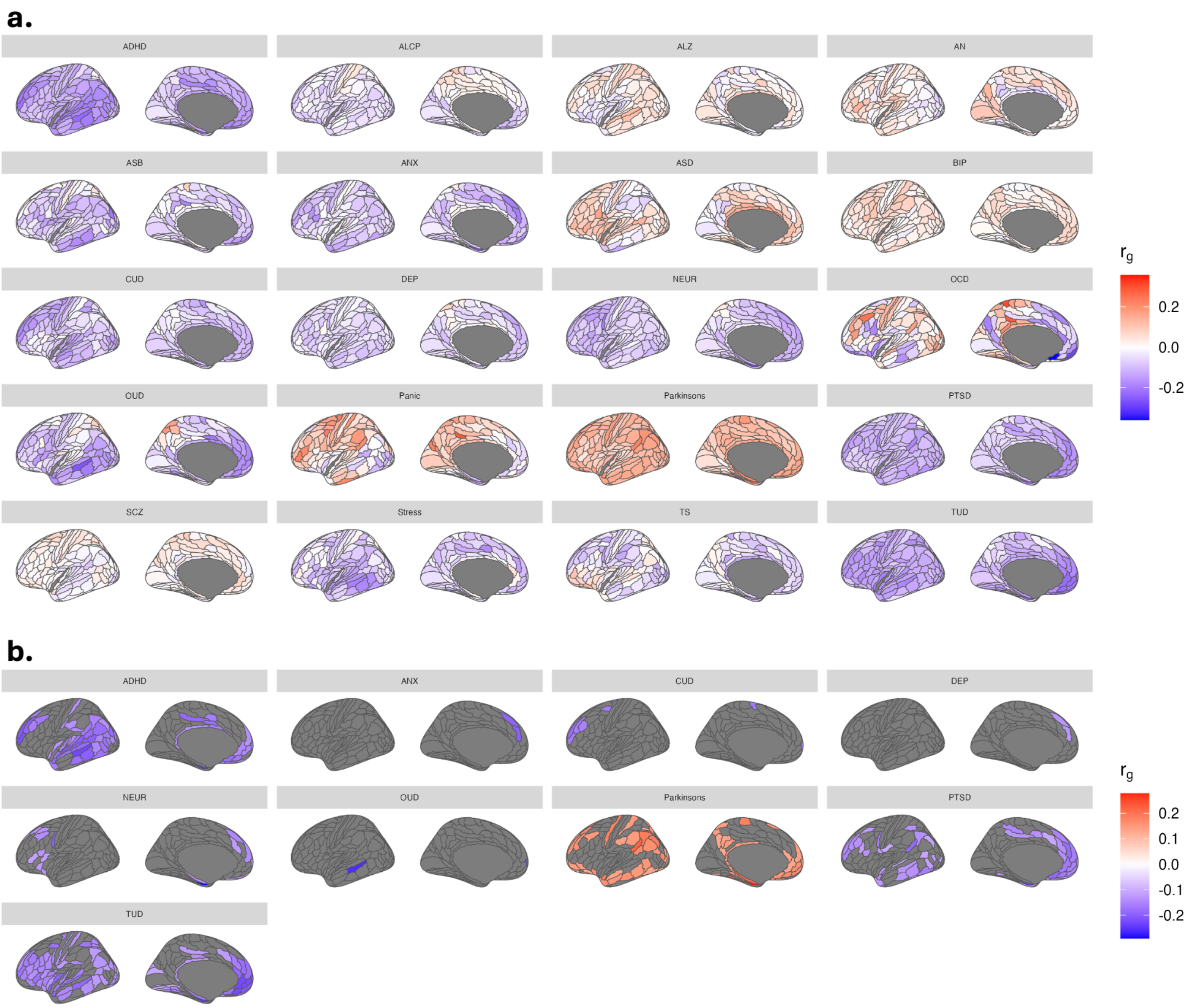


**Supplementary Figure S16. Genetic correlations between regional cortical expansion and 20 disorders.** Genetic correlations were evaluated between 177 bilateral regions and 20 neuropsychiatric disorders. **a)** Unthresholded regional results are presented. **b)** Results are thresholded by Bonferroni correction across regions based on the number of independent tests (P = 0.05/51 = 9.8e-4). ADHD: attention-deficit/hyperactivity disorder; ALCP: problematic alcohol use; ALZ: Alzheimer's disease; AN: anorexia nervosa; ANX: anxiety; ASB: antisocial behaviors; ASD: autism spectrum disorder; BIP: bipolar disorder; CUD: cannabis use disorder; DEP: depression; NEUR: neuroticism; OCD: obsessive-compulsive disorder; OUD: opioid use disorder; Panic: panic disorder; Parkinsons: Parkinson's disease; PTSD: post-traumatic stress disorder; SCZ: schizophrenia; Stress: stress-related disorders; TS: Tourette's syndrome; TUD: tobacco use disorder.

We correlated the resulting maps with each other to identify spatial similarity in the shared genetic architecture between each disorder and cortical expansion (**Fig. S17a**). Applying hierarchical clustering to this correlation matrix, we found that most externalizing disorders tended to cluster together (ADHD, ALCP, CUD, OUD, TUD) **(Fig. S17b**). Several internalizing disorders (OCD, DEP, ANX, NEUR) also clustered together, and the two neurodegenerative disorders in this study, Alzheimer's and Parkinson's, clustered together.

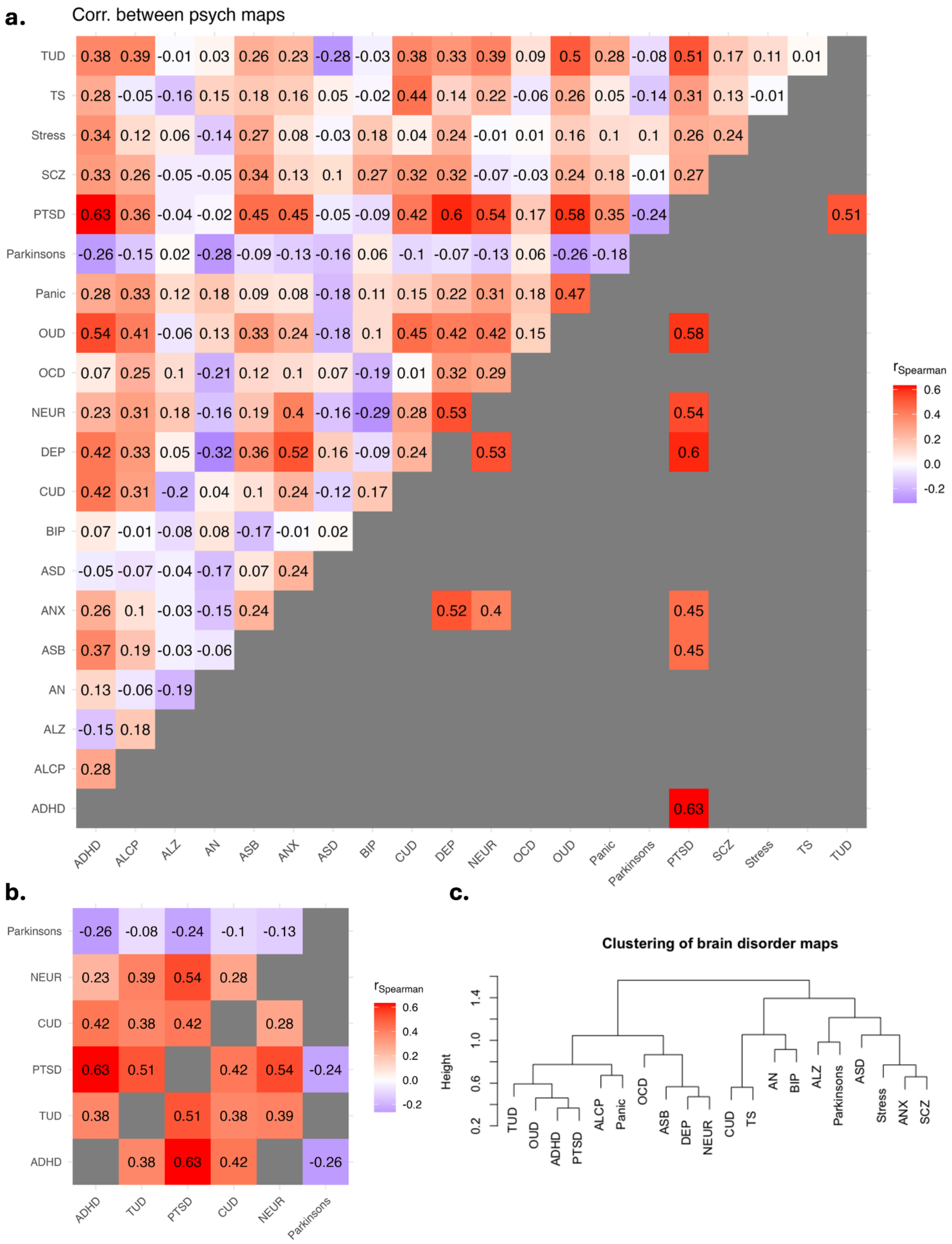


**Supplementary Figure S17. Similarity between regional maps of the correlation between cortical expansion and external disorders.** Spatial correlation between unthresholded maps of the correlations between regional cortical expansion and neuropsychiatric disorders, where the upper triangle shows all correlations, and the lower triangle shows significant correlations derived from spin tests for **a)** maps with significant global correlations, and **b)** all 20 neuropsychiatric traits. For **a)**, $P = 0.05 / 15$ pairwise tests = 3.33e-3. For **b)**, $P = 0.05 / 15$ pairwise tests = 3.33e-3. **c)** Hierarchical clustering of the correlation matrix in **a)** of unthresholded neuropsychiatric maps. This clustering process groups the disorders into groups based on having similar regional genetic relationships with cortical expansion.

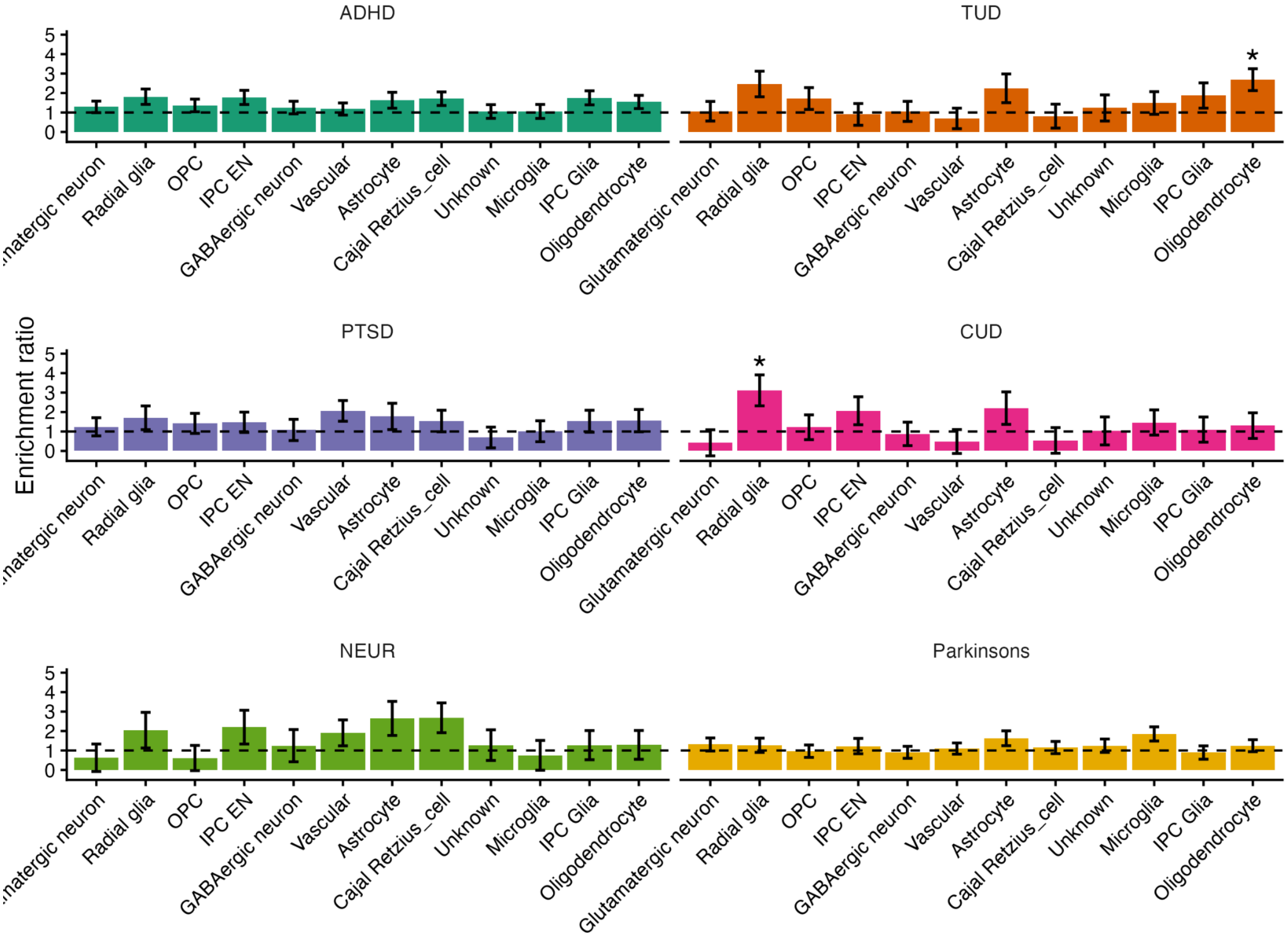


**Supplementary Figure S18. Enrichment of covariance between cortical expansion and six external traits via Stratified Genomic SEM.** The error bars represent the standard error of the enrichment ratio. Significance was determined within each disorder based on a Bonferroni correction for the number of annotations (P=0.05/12=0.004). **Cellular abbreviations:** IPC Glia: intermediate progenitor cell for glia; IPC EN: intermediate progenitor cell for glutamatergic excitatory neurons; OPC: oligodendrocyte precursor cell.

***S6. External traits MiXeR results***

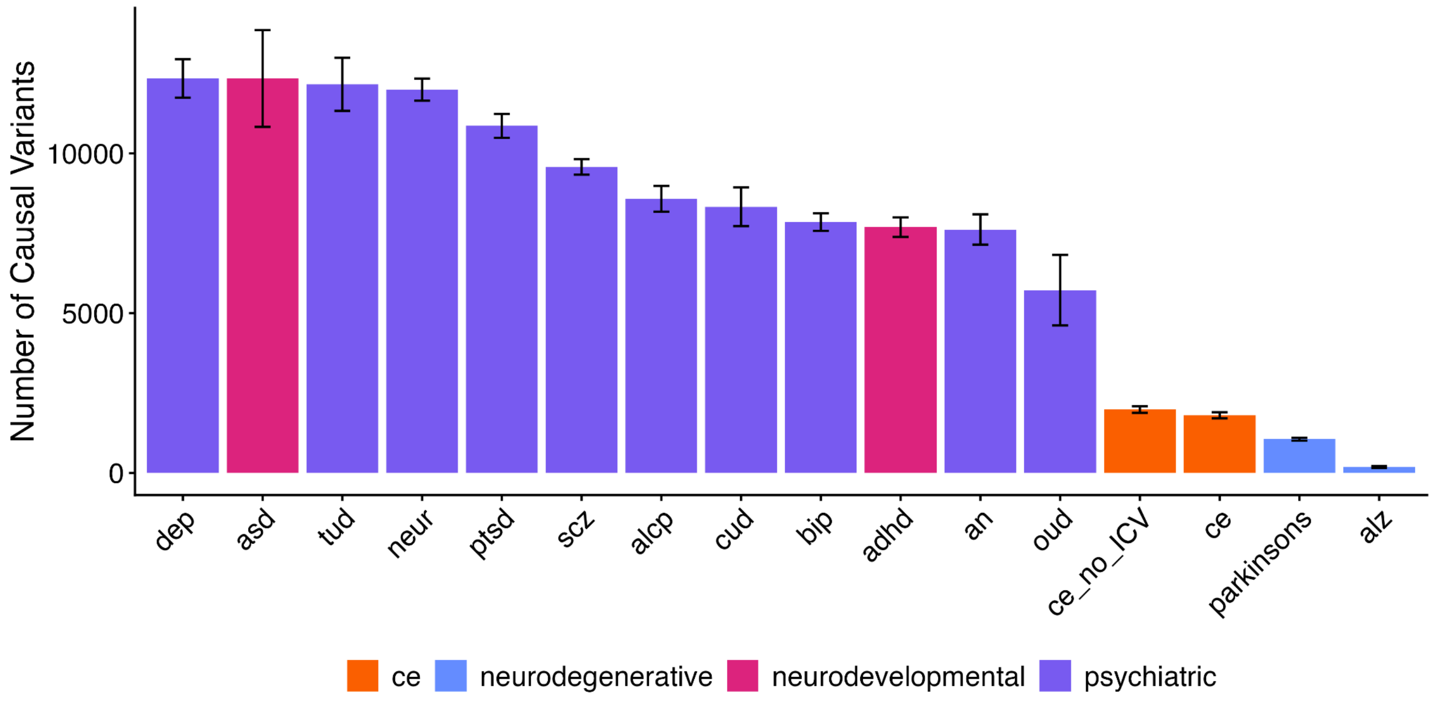


**Supplementary Figure S19. Univariate MiXeR results for neuropsychiatric traits and cortical expansion.** Univariate MiXeR estimates of the polygenicity of the cortical expansion factor and numerous other disorders, where error bars reflect the standard deviation of the estimate across 20 random iterations of MiXeR. The scale represents the estimated number of causal variants per trait at 90% SNP heritability. Neuropsychiatric disorders were broadly classified as neurodegenerative, neurodevelopmental, or psychiatric. The disorders are ordered by polygenicity (high to low).

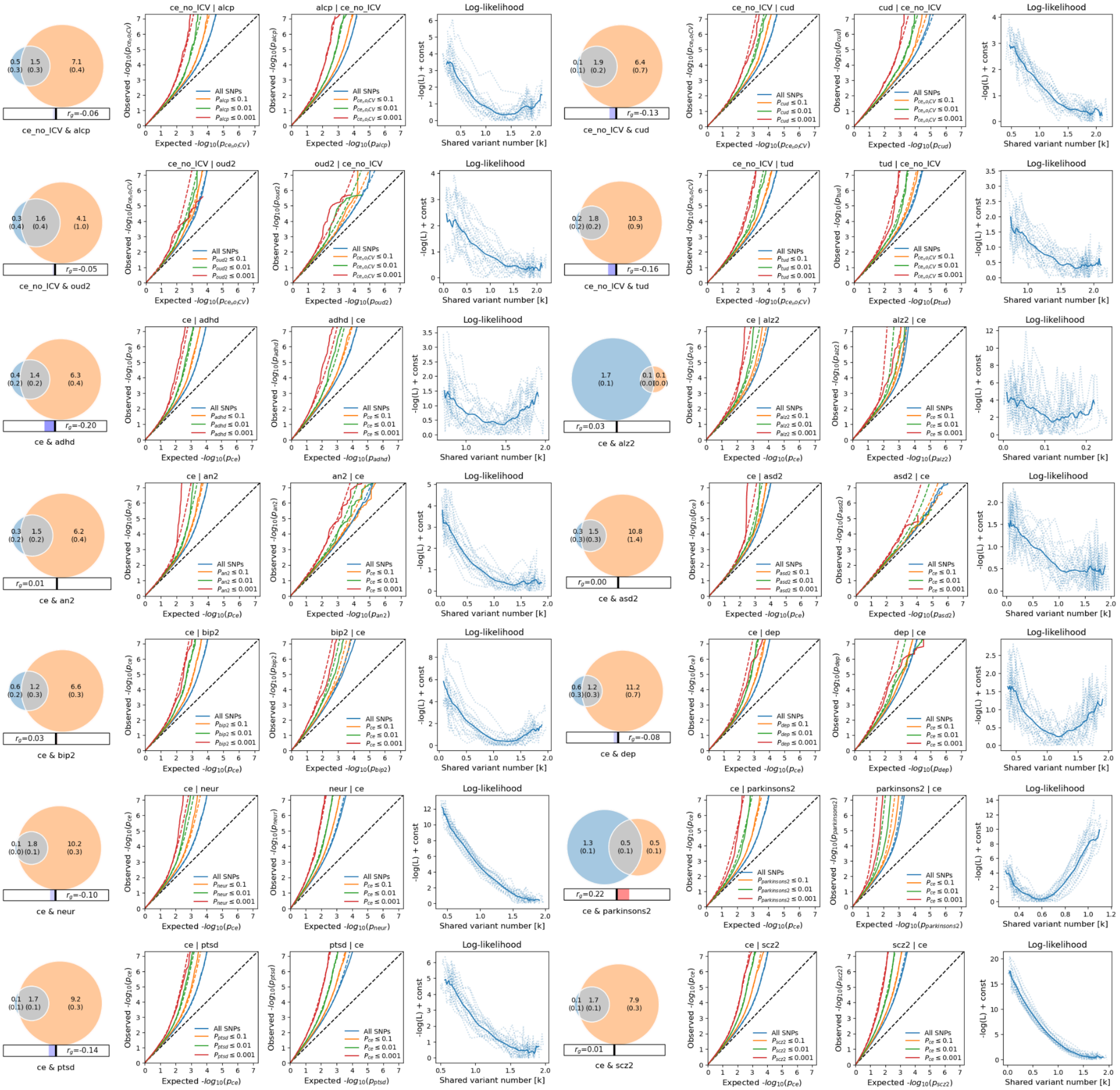


**Supplementary Figure S20. Bivariate MiXeR conditional QQ plots and log-likelihood fit for cortical expansion (CE)/disorder overlap.** For each neuropsychiatric trait with valid univariate MiXeR results, its genetic overlap (number of causal variants at 90% SNP heritability) with cortical expansion is shown as a Venn diagram. In addition, conditional QQ plots show the signal of cortical expansion GWAS stratified by the neuropsychiatric trait, and the signal of the neuropsychiatric GWAS stratified by cortical expansion signal. Finally, log-likelihood plots demonstrate the fit of the model across 20 random iterations, each selecting a random subset of 600,000 SNPs. For traits that are considered sensitive (ALCP, CUD, OUD, TUD), ICV was excluded from the CE structural equation model.
